# Structure and Mechanism of Hedgehog Acyl Transferase

**DOI:** 10.1101/2021.07.08.451580

**Authors:** Claire E. Coupland, Sebastian A. Andrei, T. Bertie Ansell, Loic Carrique, Pramod Kumar, Lea Sefer, Rebekka A Schwab, Eamon F.X. Byrne, Els Pardon, Jan Steyaert, Anthony I. Magee, Thomas. Lanyon-Hogg, Mark S. P. Sansom, Edward W. Tate, Christian Siebold

## Abstract

The iconic Sonic Hedgehog (SHH) morphogen pathway is a fundamental orchestrator of embryonic development and stem cell maintenance, and is implicated in cancers in various organs. A key step in signalling is transfer of a palmitate group to the N-terminal cysteine residue of SHH, catalysed by the multi-pass transmembrane enzyme Hedgehog acyltransferase (HHAT) resident in the endoplasmic reticulum (ER). Here, we present the high-resolution cryo-EM structure of HHAT bound to substrate analogue palmityl-coenzyme A and a SHH mimetic megabody. Surprisingly, we identified a heme group bound to an HHAT cysteine residue and show that this modification is essential for HHAT structure and function. A structure of HHAT bound to potent small molecule inhibitor IMP-1575 revealed conformational changes in the active site which occlude substrate binding. Our multidisciplinary analysis provides a detailed view of the novel mechanism by which HHAT adapts the membrane environment to transfer a long chain fatty acid across the ER membrane from cytosolic acyl-CoA to a luminal protein substrate. This structure of a member of the protein-substrate membrane-bound O-acyltransferase (MBOAT) superfamily provides a blueprint for other protein substrate MBOATs, such as WNT morphogen acyltransferase Porcupine and ghrelin *O*-acyltransferase GOAT, and a template for future drug discovery.

## INTRODUCTION

Hedgehog (HH) signalling is essential for human embryogenesis and stem cell regulation, and mutations in pathway members often cause congenital diseases, such as holoprosencephaly (Briscoe and Therond, 2013; Ingham et al.). Aberrant HH signalling is linked to the formation of numerous cancers, and it has been estimated that around 25% of all fatal cancers show deregulated HH activation, making the pathway an important target for cancer drug discovery (Chahal et al., 2018; Wu et al., 2017). Post-translational protein lipidation is integral to HH signalling; the HH morphogen Sonic hedgehog (SHH) undergoes a highly unusual and unique maturation process in which a 45 kDa preprotein undergoes autocleavage and concomitant attachment of a cholesterol moiety at the new C-terminus (Bumcrot et al., 1995; Porter et al., 1995; Porter et al., 1996). Following cleavage of a signal peptide, a palmitoyl (C16:0) acyl group is transferred to the new N-terminal cysteine (Cys24) in the endoplasmic reticulum (ER) (Pepinsky et al., 1998) to yield the mature 19 kDa, dually-lipidated SHH signalling domain. Hedgehog acyltransferase (HHAT) catalyses the irreversible transfer of a palmitoyl group from palmitoyl-coenzyme A (Palm-CoA) to the N-terminal nitrogen of SHH-Cys24 (Buglino and Resh, 2008; Chamoun et al., 2001), which is the rate-limiting step in HH ligand production. Mechanistically, it is currently unknown whether HHAT directly *N*-acylates HH proteins, or whether this occurs via S-acylation of Cys24 followed by *S*,*N*-acyl shift to yield the *N*-acylated product. Lipidation of SHH is required for interaction with its receptor Patched (PTCH1) at the signal receiving cell surface (Kong et al., 2019), whereby the N-terminal palmitoyl group binds to PTCH1 and relieves inhibition of the G protein-coupled receptor Smoothened (SMO) to activate downstream HH signalling (Qi et al., 2018; Tukachinsky et al., 2016). Accordingly, HHAT knockout leads to a loss-of-function HH phenotype (Chen et al., 2004) and HHAT has been identified as a potential cancer drug target (Konitsiotis et al., 2014; Matevossian and Resh, 2015a; Petrova et al., 2013; Rodgers et al., 2016).

HHAT is a member of the membrane-bound *O*-acyltransferase (MBOAT) superfamily (Hofmann, 2000). The majority of MBOATs catalyse lipid chain transfer to small molecule lipid substrates, however HHAT, alongside porcupine acyltransferase (PORCN) and ghrelin acyltransferase (GOAT), acylates protein substrates. PORCN catalyses *O*-palmitoleoylation of Wnt in development (Herr and Basler, 2012), while GOAT catalyses ghrelin *O*-octanoylation in appetite regulation (Gutierrez et al., 2008; Yang et al., 2008). To date, structures have been determined for three MBOATs: bacterial DltB (Ma et al., 2018), and human ACAT (Long et al., 2020; Qian et al., 2020) and DGAT (Sui et al., 2020; Wang et al., 2020). These share a common 8 transmembrane (TM) helix core domain that contains the active site including a conserved histidine (Hofmann, 2000) and in the cases of AGAT and DGAT the acyl co-enzyme A substrate binding site. Topological studies have suggested that HHAT contains a similar domain (Konitsiotis et al., 2015; Matevossian and Resh, 2015b). However, in the absence of any structural information on HHAT, the mode of palmitoyl transfer to its protein substrate SHH remains elusive. Here, we present a high resolution cryo-EM structure of HHAT in complex with the non-hydrolysable Palm-CoA analogue palmityl-CoA (nhPalm-CoA) and an inhibitory megabody. Our structural analysis reveals a novel cavity for SHH binding on the ER luminal face that leads to the active site. Surprisingly, we identified a heme molecule bound proximal to the palmitoyl-CoA binding site that plays a crucial functional role. A further HHAT structure with a potent small molecule inhibitor IMP-1575 bound identified significant conformational changes in the active site, which block palmitoyl-CoA access to SHH. Our results clarify the structural mechanism by which an MBOAT family member can catalyse acylation of a protein substrate whilst exchanging a lipid substrate across the ER membrane, and provide a template for future development of a pharmacological inhibitor.

## RESULTS

### Overall structure of human HHAT

To structurally and functionally characterise HHAT, we raised camelid antibodies (nanobodies, NBs) against full-length human HHAT and isolated two NBs (NB169 and NB177) that bound to HHAT expressed in cells and in biolayer interferometry using purified HHAT with low nanomolar affinity (Fig. S1). To overcome the technical hurdles of size and/or preferential orientation problems for cryo-EM structure analysis, we inserted the nanobodies into the *E. coli* K12 glucosidase YgjK scaffold resulting in ∼100 kDa “megabodies” (Uchanski et al., 2021) (megabodies MB169, MB177). We reconstituted the complex of human HHAT (incubated with nhPalm-CoA) and MB177 in detergent micelles and determined the cryo-EM complex structure to 2.69 Å resolution using a localised refinement strategy (Fig. S2). The high quality cryo-EM map allowed building of almost all sidechains of HHAT and MB177 as well as nhPalm-CoA (Fig. S2A).

Our cryo-EM analysis revealed that HHAT forms a 1:1 complex with MB177 (Fig. 1A**, Movie S1**), which is consistent with multiangle light scattering analysis (SEC-MALS) carried out in solution (Fig. S1F). HHAT is composed of 12 TM helices with both termini located on the cytosolic side of the ER membrane (Fig. 1B, Fig. S3). The 8-TM helix core of HHAT resembles that of the other MBOAT superfamily members (Fig. 1C, coloured in blue). However, the HHAT N- and C-terminal regions are unique and distinct from other MBOAT structures (Fig. 1C, coloured in orange and green, Fig. S4). These rearrangements distort the dimerization mode observed in DGAT and ACAT (which mediate attachment of fatty acids to lipids), thereby occluding the lateral gate into the lipid bilayer that is proposed to allow lipid substrate entry (Long et al., 2020; Qian et al., 2020; Sui et al., 2020; Wang et al., 2020). Instead, the rearrangement of the N- and C-terminal helices results in the formation of a distinct pocket that is solvent accessible from the cytosolic side.

**Fig. 1.**
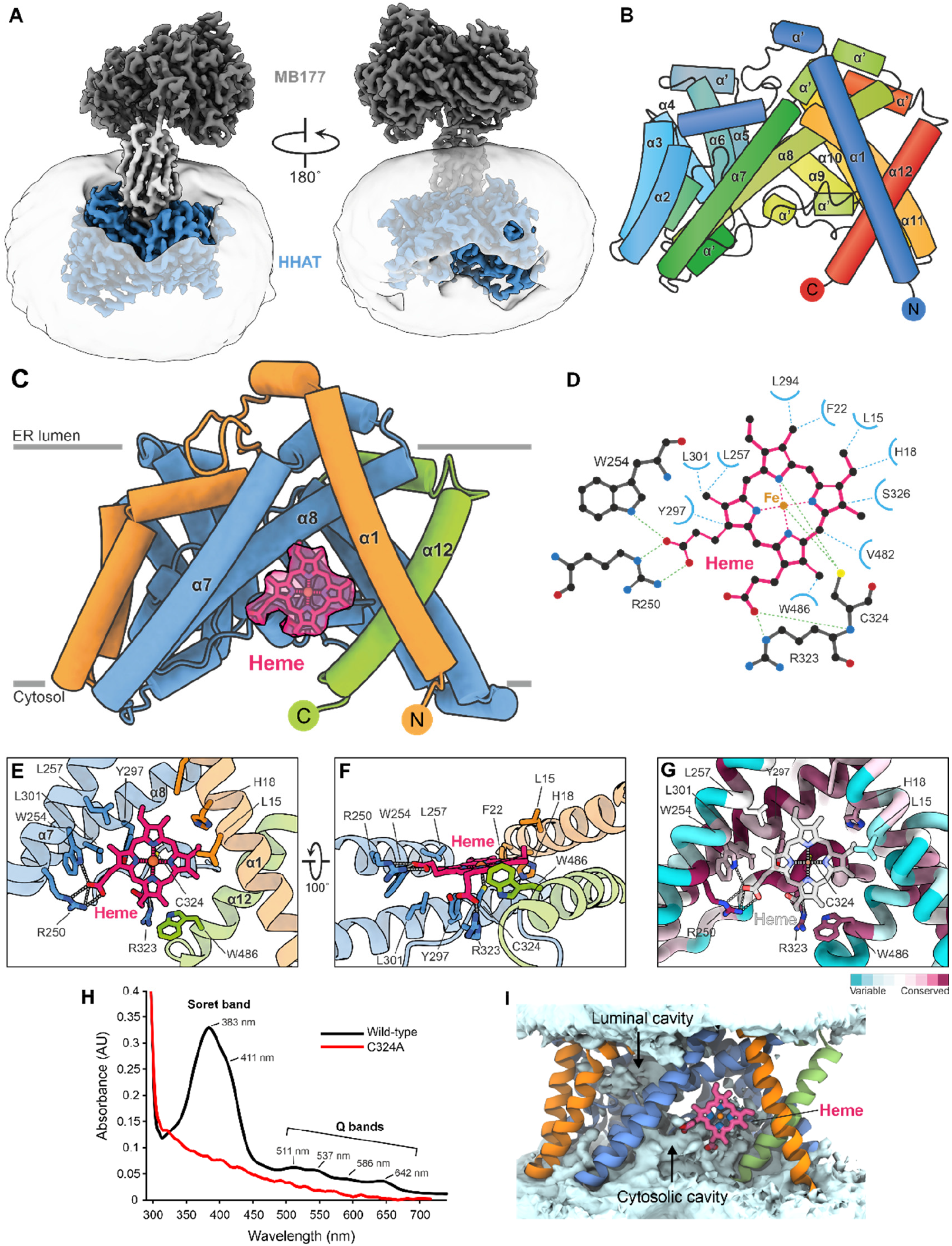
Structure of human HHAT. **(A)** Cryo-EM map of the HHAT-megabody 177 complex at 2.69 Å resolution. HHAT is coloured in blue, the megabody scaffold in dark grey and the nanobody in light grey. The detergent micelle is depicted. **(B)** Schematic of HHAT in rainbow colouring (blue: N-terminus, red: C-terminus). **(C)** Cartoon representation of human HHAT. The structurally conserved MBOAT core is coloured in blue. The variable N-(orange) and C-(green) termini are highlighted. The heme-B group is depicted in magenta stick representation with its cryo-EM map shown. **(D)** Schematic of the heme-B interactions with HHAT residues. **(E-G)** Close-up views of the HHAT heme-B binding site. **F** and **G** are coloured as in **C**. The sequence conservation (purple: conserved to green: variable) is mapped onto the structure in **G**. **(H)** UV/VIS spectra of HHAT wildtype (black) and HHAT-C324A (red). Wildtype HHAT shows the typical spectrum for a penta-coordinate thiolate Fe(III) heme with the split Soret band (at 383 nm and 411 nm) and the four Q bands (I: 642 nm, II: 586 nm, III:537 nm and IV: 511 nm). **(I)** Time-averaged solvent density (light blue surface) across 5x 200 ns atomistic simulations of HHAT coloured as in **C**.

### HHAT is a heme protein

Surprisingly, we observed strong additional density in this solvent accessible pocket that we identified as a heme-B molecule (Fig. 1C). The major interaction is formed by the sidechain of Cys324 that axially coordinates the Fe(III) ion of the heme-iron complex. Heme binding is supported by various, conserved hydrophobic interactions (e.g. Leu257, Trp254 and Trp486) and two salt bridges between HHAT residues Arg250 and Arg323 with the two carboxylate groups of the heme porphyrin ring. (Fig. 1D-F). The heme binding site is highly conserved across HHAT homologues (Fig. 1G and Fig. S3). To confirm the presence of an HHAT-bound heme moiety, we carried out UV/VIS spectroscopy (Fig. 1H) and observed a spectrum typical of a cysteine-coordinated heme protein characterised by a broad Soret band around 380 nm, followed by several weaker absorptions (Q bands) at higher wavelengths (from 500 to 650 nm) (Kuhl et al., 2013; Uppal et al., 2016) HHAT-C324A, in which the cysteine that coordinates the heme iron is mutated to alanine, lacks the Soret and Q bands, further supporting the presence of an HHAT-heme complex in the structure. Cys324 was previously shown to be important for HHAT folding and suggested to be post-translationally modified in addition to other cysteine residues (Konitsiotis et al., 2015). The strong heme density on Cys324 is clearly distinct from the lipid-like attachments we observe for multiple cysteines in the HHAT structure (Fig. S5) with partial density consistent with reversible long chain *S*-acylation.

The rearrangement of the termini forms a cytoplasmic cavity with the heme molecule at its apex, which is solvent accessible from the cytosolic side of the membrane in atomistic simulations (Fig. 1I, Fig S6). While visualisation of the protein structure alone might suggest that the heme moiety would be embedded within the hydrophobic bilayer core, MD simulations indicate that it is situated at the membrane-solvent interface, with the heme carboxyl groups puncturing the wetted cytoplasmic cavity. Thus, heme binding could be regulated by both membrane and cytoplasmic components.

### Mode of palmitoyl-CoA binding

We determined HHAT in complex with a non-hydrolysable Palm-CoA analogue lacking the thioester carbonyl (nhPalm-CoA), which is bound within the HHAT core domain (Fig. 2A and 2B**, Movie S1**). The entire 16:0-CoA molecule is visible in the cryo-EM map. nhPalm-CoA binding occludes a continuous solvent cavity through HHAT that extends from the cytoplasmic cavity towards the luminal face (Fig. S6). The CoA adenine ring protrudes into the solvent at the cytosolic site, stacking between Ile352 and Gln358, and the N1 and N6 nitrogen atoms form hydrogen bonds with the amide nitrogen of Ser357 and the carbonyl oxygen of Gly355, respectively (Fig. 2C and 2D). The CoA phosphate groups are coordinated by a metal ion, built as a magnesium ion based on coordination geometry (Harding, 2001), and several hydrogen bonds mediated by Arg336 and Ser357. The Palm-CoA moiety is shifted along the membrane normal in the HHAT binding site compared to its average simulated position in a membrane bilayer (Stix et al., 2020). This locates the Palm-CoA thioester in the core of the HHAT cavity close to the proposed reaction center residues His379 and Asp339 (Fig. 2E) that have previously been shown to be important for catalysis of palmitoyl transfer to SHH (Buglino and Resh, 2010; Hofmann, 2000).

**Fig. 2.**
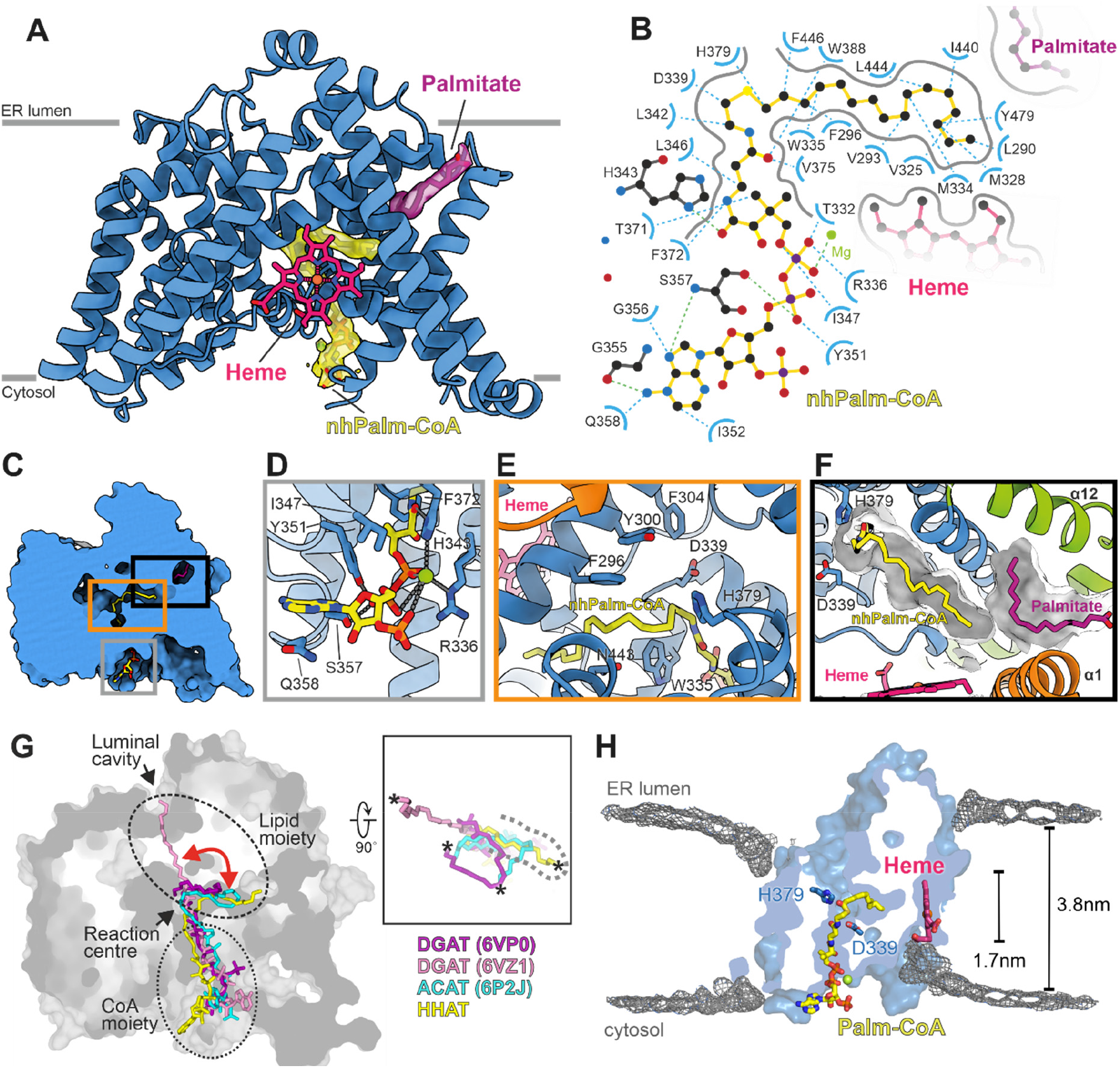
Structure of HHAT in complex with its co-factor analogue nhPalm-CoA. **(A)** Cartoon representation of HHAT with bound nhPalm-CoA (yellow) and a palmitate molecule (purple) that forms part of the Palm-CoA binding site. The 2.69 Å resolution cryo-EM map for selected molecules is shown. **(B)** Detailed interactions of the Palm-CoA binding site. Hydrogen bonds are depicted in blue, the magnesium ion is in green. **(C-F),** Close-up views of the Palm-CoA binding site. **(G)** Structural superposition of HHAT with the DGAT-oleoyl-CoA complexes (PDB ID. 6VZ1 and 6VP0, respectively) and the ACAT-oleoyl-CoA (PDB ID. 6P2J) complexes, with only the acyl-CoA moieties depicted. The HHAT solvent accessible surface is shown. Lipid and Co-A moieties of the acyl-CoA substrates are circled. The red arrow indicates the conformational flexibility of the CoA substrate acyl chains. The close-up in the right panel highlights the different conformations of the acyl chains viewed from the ER luminal side. The HHAT palmitate binding pocket is marked with a dotted line. The terminal carbon atom of the acyl-CoA substrate is depicted with an asterisk for clarity. **(H)** Time averaged density of lipid phosphate beads (grey isomesh) across 10x 15 μs CG simulations of HHAT overlayed with the atomistic HHAT structure, coloured as in **A**. HHAT is depicted as a cut surface through the centre of the protein to indicate the respective positions of the deformations.

The tail of the nhPalm-CoA 16:0 chain inserts into a binding pocket formed by rearrangement of the HHAT N- and C-terminal helices (Fig. 2E and 2F). The HHAT palmitoyl binding pocket is unique and is positioned to occlude the dimer interface observed in lipid MBOATs, mainly because of rearrangement of the N- and C-terminal helices (Fig. S4). The palmitoyl moiety forms mainly hydrophobic interactions with residues located in the HHAT core, however we also observe additional kinked lipid density that forms part of the palmitoyl substrate binding site, modelled as an additional palmitate moiety (Fig. 2F). This novel arrangement of the termini restricts the acyl chain length, to a maximum of 16 carbons, and provides a rationale for previous analyses of SHH expression revealing acylation of the SHH N-terminus predominantly by shorter acyl chains (e.g. myristate and myristoleate) in addition to palmitate, and to a far lesser extent by longer lipids (Long et al., 2015; Pepinsky et al., 1998). In contrast the acyl binding pocket of lipid MBOATs appears to be shallower and less defined, and the acyl chains show a high degree of conformational flexibility, even extending towards the luminal side (Fig. 2G).

An intriguing question arising from HHAT function is how it may reduce the energetic barrier for palmitoyl transfer across the membrane. Coarse grained (CG) MD simulations of HHAT in a model ER membrane revealed substantial bilayer deformation around two regions in opposing leaflets formed by Leu321-Thr327 (heme binding site) and around α helices 5 and 9 (Fig. 2H, Fig. S6G and S6H). These bilayer deformations funnel towards the reaction center with a net reduction in bilayer width of up to ∼50 % (1.7 nm) compared to the width at extended distances from the protein (3.8 nm). Thus, HHAT-induced bilayer thinning may reduce energetic barriers in the catalytic cycle and/or aid entry/exit of bound ligands.

### Structure of HHAT bound to small-molecule inhibitor IMP-1575

Given the importance of HHAT as a potential target to inhibit SHH signalling in disease, we determined the structure of HHAT bound to IMP-1575, the most potent HHAT small molecule inhibitor reported to date (Lanyon-Hogg et al., 2021) (Fig. 3A). IMP-1575 binds in the middle of the reaction centre, forming a hydrogen bond to the catalytic HHAT His379 and thus occluding Palm-CoA binding (Fig. 3B and 3C). Our previous computational analysis suggested that the stereogenic centre of IMP-1575 has (*R*) absolute stereochemistry (Lanyon-Hogg et al., 2021), and this stereochemistry is confirmed in the cryo-EM structure. IMP-1575 binding leads to a rearrangement of the reaction centre when compared to the HHAT-nhPalm-CoA complex, affecting the sidechain conformations of the active site Asp339 as well as Asn443 and Trp335 (Fig. 3C). Trp335 is situated in close proximity to the Palm-CoA thioester bond and exhibits a dramatic conformational switch between the two structures such that the side chain is rotated inwards towards the Palm-CoA binding pocket when inhibitor is bound (**Movie S1**). This is consistent with kinetic studies of HHAT inhibition by IMP-1575 and rationalises the mechanism of Palm-CoA competitive inhibition by IMP-1575 (Lanyon-Hogg et al., 2021).

**Fig. 3.**
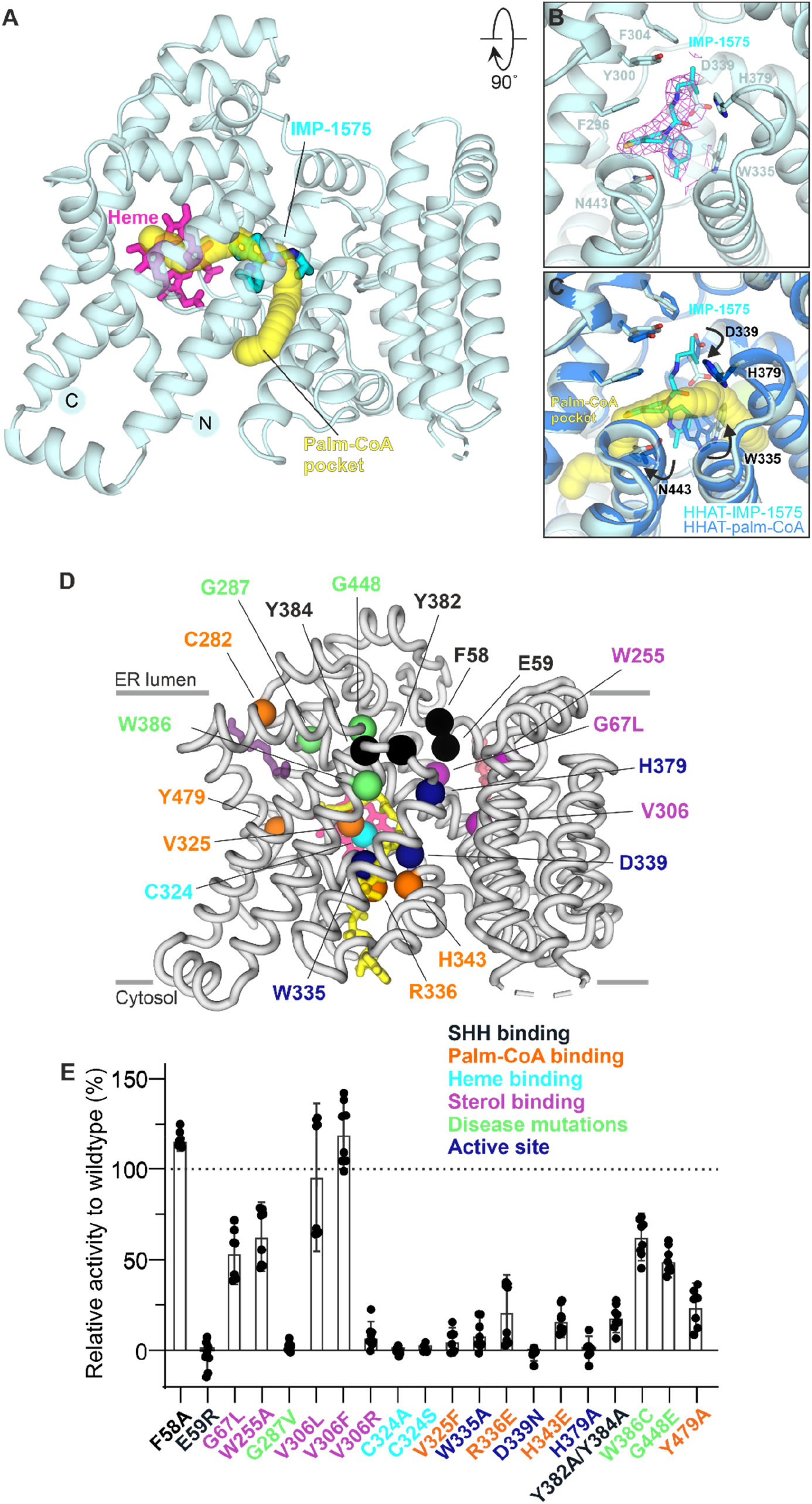
Structure of HHAT in complex with inhibitor IMP-1575. **(A)** Structure of HHAT in complex with the inhibitor IMP-1575. The Palm-CoA binding pocket calculated with program CAVER is depicted in yellow, the bound heme moiety is in magenta and IMP-1575 is highlighted in stick representation. View is rotated 90° around the x axis compared to Fig. 1C. **(B)** Close-up view of the IMP-1575 binding site. IMP-1575 binds in the active site in close proximity to the catalytic His379. The cryo-EM map of the HHAT-IMP-1575 complex is shown as pink chicken wire. **(C)** Superposition of the HHAT-nhPalm-CoA (marine) and -IMP-1575 complexes (cyan). The Palm-CoA binding pocket is shown in yellow. The arrows indicate the conformational changes induced by inhibitor binding. IMP-1575 occludes Palm-CoA binding. Orientation is as in **B**. **(D, E)** Structure-function analysis of HHAT mutants. A ribbon representation of HHAT with residues mutated in spheres is shown in **D**. nhPalm-CoA (yellow), heme (pink), palmitate (dark purple) and cholesterol (red) are shown in sticks representation. Activities of HHAT mutants relative to wildtype HHAT, measured by Acyl-cLIP assay. N = 2 (quadruplicates), mean ± SEM, black dots are shown in **E** and represent replicates within biological replicates.

To probe Trp335 dynamics, we performed atomistic MD simulations of HHAT. When the Palm-CoA binding site was unoccupied (starting from both the Palm-CoA or IMP-1575 bound structures), the Trp335 indole ring moved towards Phe372 to close the Palm-CoA tunnel (Fig. S7A-G). This conformation poses a hydrophobic restriction in the Palm-CoA binding pocket. In contrast, when Palm-CoA was bound, Trp335 remained in an upright conformation. Thus, water permeation through HHAT is prevented by either Palm-CoA or Trp355 movement. In simulations, we also observe dynamic behaviour of the Asp339 side chain, which either faces towards Gly341/Leu342 (‘pocket facing’) or toward the solvated catalytic core (‘His379 facing’). Palm-CoA binding restricts Asp339 to the ‘pocket facing’ conformation due to steric occlusion of the ‘His379 facing’ conformation (Fig. S7H and S7I). We estimated the pKa of Asp339/His379 across the simulations (Fig. S7J), yielding a mean pKa for Asp339 of ∼5.6 and for His379 of ∼4.6. The pKa values are shifted due to a large number of proximal hydrophobic residues and, crucially, the donor/acceptor pairing is reversed suggesting these residues do not function in a proton shuttle mechanism. The ability of His379 to form an H-bond with IMP-1575, along with proximity to the Palm-CoA reaction centre and the shifted pKa, may suggest a role acting as a H-bond donor in activation of the Palm-CoA carbonyl in catalysis.

### Structure-activity analysis to identify key features for HHAT function

In order to investigate the role of the observed structural features in catalysis, we generated and purified a series of structure-guided HHAT mutants. These were evaluated for enzymatic activity by monitoring palmitoylation of a FAM-labelled substrate peptide (**CGPGRGFGKR**(K-FAM)G-CONH_2_) in real time using the Acyl-cLIP assay (Lanyon-Hogg et al., 2019) (Fig. 3D and 3E). Mutation of the canonical MBOAT active site residues Asp339 and His379 (D339N and H379A) abolishes activity, as does mutation of heme-binding Cys324 (C324A & C324S). The distance (∼19 Å) and lack of a clear connection between the heme Fe ion and the reaction centre argues against a direct role in catalysis and supports a suggested role of Cys324 in HHAT folding (Konitsiotis et al., 2015). Mutation of Trp335 to alanine (W335A) also reduced HHAT activity, consistent with its proposed role as a gate-keeper residue of the palm-CoA pocket. Modification of the phosphate/magnesium binding site for nhPalm-CoA (R336E and H343E) at the cytosolic site completely abrogated activity, similar to V325F, which was designed to block the Palm-CoA fatty acid binding tunnel. This is also seen for Y479A which interferes with shielding the acyl chain of the nhPalm-CoA from direct contact with the heme group. This confirms the observed Palm-CoA binding site in our structure and as shows that proper Palm-CoA binding is essential for HHAT function, as expected.

On the luminal face, Phe58 and Glu59 are hypothesised to interact with the incoming SHH substrate. Whereas modification of Phe58 (F58A) did not markedly affect activity, a charge reversal at E59R leads to a complete loss of activity, presumably due to repulsion of the positively charged character of the SHH N-terminal peptide substrate. Moreover, alanine exchange of the tyrosine residues located above the reaction centre (Tyr382 and Tyr384), and thus likely interacting with SHH, significantly reduces HHAT activity. We also investigated the importance of an observed sterol binding site that is located adjacent to luminal SHH-binding cavity, and which has Val306 at its centre and is lined by Gly67 and Trp255 (Fig. S9). Mutation of Val306 to hydrophobic residues (V306L and V306F) is tolerated, but introduction of a positive charge (V306R) severely reduces enzymatic activity. G67L and W255A show some effects, however still retain some 50% activity, presumably either by maintaining packing or allowing a sterol molecule to bind. Taken together, our structure-guided mutagenesis analysis supports the structure features observed in the cryo-EM map and identifies key regions important for HHAT function.

Finally, we tested several known disease-associated HHAT mutations.The G287V mutation was reported to abrogate catalytic activity and induce Syndromic 46,XY Disorder of Sex Development (DSD) (Callier et al., 2014). Our results confirm this, and our structure suggests this mutation would severely distort the packing of helix α8 and the N-terminal helix α1, disrupting the heme binding site and potentially rearranging the palmitoyl binding tunnel and/or reaction centre. Mutation W386C has been implicated in intellectual disability (Agha et al., 2014) and shows a minor reduction in enzymatic activity according to our data. Our structure has clear density for a palmitic acid moiety in this area, and so a mutation to cysteine may lead to palmitoylation at this position. The G448E mutation was identified in a melanoma cell-line and may play a role in cancer development (Kawakami et al., 2001). Our structure shows it is located at the rim of the luminal access cavity for SHH. Even though our enzymatic data shows only a modest decrease in activity, we hypothesise that its effect could be more profound on palmitoylation of full-length SHH. A glutamate residue at this position would protrude straight into the access channel, potentially blocking the larger protein from entering the HHAT active site. We also attempted to express the L257P mutant, which causes microcephaly (Abdel-Salam et al., 2019), but expression levels were too low to isolate the protein in useful quantities (data not shown). Leu257 is located in helix α7 and forms direct hydrophobic contacts to the heme group (Fig. 1F). We hypothesise that mutation to proline would not only be disruptive to heme binding, but also cause a helix kink that could disrupt the core MBOAT fold, leading to misfolded protein. In summary, our data provides a mechanistic rationale for observed disease-linked mutations in HHAT.

### A novel luminal access cavity for SHH

Our structural analyses show that the lipid substrate entry site in the membrane in other MBOATs is occluded by rearrangement of the HHAT termini (Fig. S4). How does SHH, a hydrophilic protein substrate, access the HHAT active site? Our structure reveals a distinct luminal cavity arrowing towards the catalytic His379 in the reaction centre, where the major HHAT interaction loop of MB177 (CDR3 loop) is located, suggesting an access path for SHH (Fig. 4A, Fig. S8). This cavity is highly conserved in all HHAT family members (Fig. 4B) and forms part of a larger acidic interface spanning the luminal surface (Fig. 4C). This negative patch complements the positively charged N-terminus of human SHH (C24-GPG**R**GFG**KRRH**P**KK**), the first six residues of which are essential for HHAT-mediated palmitoylation (Hardy and Resh, 2012). This N-terminal stretch is disordered in structures of SHH in complex with HHIP (Bishop et al., 2009; Bosanac et al., 2009) or CDO (McLellan et al., 2008), or apo SHH (Hall et al., 1995), and is only visible in the SHH-PTCH1 complex structure where the N-terminal SHH palmitate protrudes into the PTCH1 core (Qi et al., 2018; Rudolf et al., 2019).

**Fig. 4.**
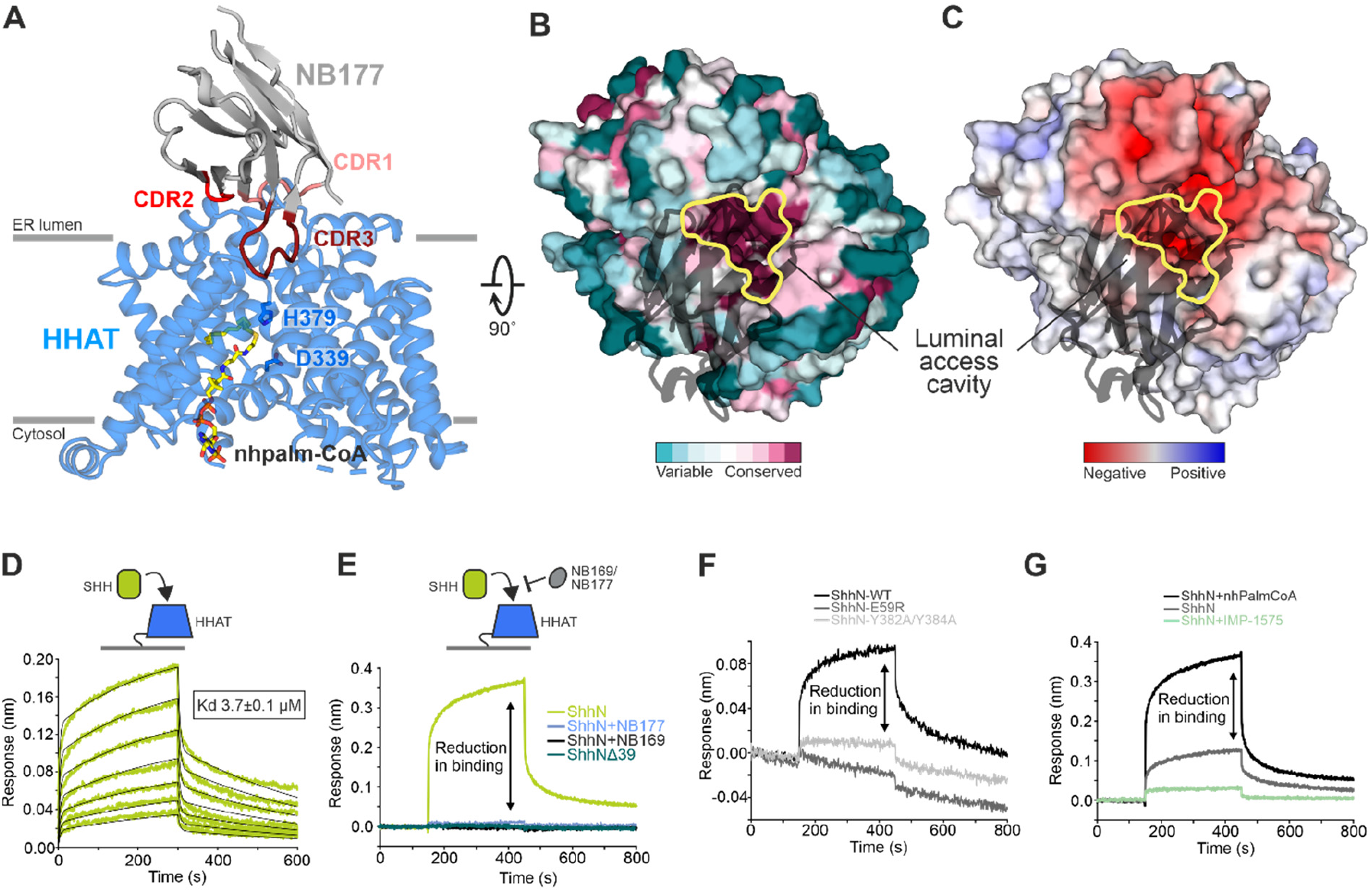
A luminal access cavity for SHH. **(A)** Cartoon representation of the HHAT-NB177 complex, HHAT is coloured in blue, NB177 in grey with the complementary determining regions (CDRs) coloured in salmon (CDR1), red (CDR2) and dark red (CDR3). Active site HHAT-residues His379 and Asp339 are highlighted in stick representation. The active site entry tunnel, which is mainly blocked by NB177-CDR3, protrudes deeply into HHAT. **(B, C)** Surface representation of HHAT with NB177 shown in black ribbon. Sequence conservation (**B**) and electrostatic potential (**C**, from ± 8kT/eV) and are mapped onto the solvent accessible HHAT surface. The active site entry tunnel is outlined in yellow. Orientation is 90 ° rotated around the y axis. **(D)** Multicycle kinetics measurement of the ShhN-HHAT interaction with real-time sensorgrams for different ShhN concentrations. **(E)** ShhN binding is abolished in a SHH construct missing the first 15 residues, or in the presence of MB169 and MB177. HHAT mutants E59R and Y382A/Y384A, located in the luminal binding cavity, cannot bind SHH **(F),** and SHH binding reduced by 50% in absence of Palm-CoA and completely abolished in the presence of inhibitor IMP-1575 (**G**).

We also determined the structure of HHAT in complex with another megabody (MB169), which shares the same epitope and CDR loop conformation, despite belonging to a different nanobody family (sequence identity of CDR regions < 28%) (Fig. S1 **and S8**). In these structures, the CDR3 loops of NB169 and NB177 penetrate the luminal cavity, binding in close proximity to the active H379 (∼5Å distance) (Fig. S8). Both nanobodies as well as corresponding megabodies are efficient inhibitors of SHH palmitoylation, inhibiting palmitoyl transfer to a SHH peptide with IC_50_ values of 1.3 nM (NB169), 0.9 nM (NB177), 7.5 nM (MB169) and 31.5 nM (MB177) (Fig. S8G). This suggests that the nanobodies and SHH compete for access to the active site via the luminal cavity. To further test this hypothesis, we set up a competition binding assay using using bio-layer interferometry (BLI). The full-length N-terminal signalling domain of SHH (ShhN) only lacking the lipid attachments bound HHAT with a Kd of 3.7 μM (Fig. 4D), whereas an SHH truncation, missing the 15 N-terminal residues essential for palmitoylation (ShhNΔ39) was not able to bind HHAT (Fig. 4E). When carrying out the same experiment in the presence of a constant concentration of 200 nM NB169 or MB177, ShhN binding was occluded, confirming that the luminal cavity of HHAT is the access route for SHH (Fig. 4E). To support our competition binding experiments, we tested HHAT mutants E59R and Y382A/Y384A, both located at the apex of the luminal cavity, in our BLI assay (Fig. 4F). Both mutants severely impaired SHH binding, in agreement with our activity assay showing no (for E59R) or reduced (for Y382A/Y384A) palmitoylation. Interestingly, the ShhN-HHAT interaction was reduced by some 50% in the absence of nhPalm-CoA and almost completely abolished in the presence of IMP-1575 (Fig. 4F), suggesting that SHH binding is dependent on Palm-CoA binding to HHAT, likely due to Palm-CoA-mediated reorganisation of Trp335 and/or Asp339.

## Discussion

Cysteine-bound heme-B iron complexes in prototypical heme proteins (such as hemoglobin,myoglobin or cytochrome P450) play key roles in oxygen storage, transfer and activation, as well as being mediators of electron transfer. We identified a heme-B bound to HHAT and show that mutation of the heme-binding cysteine results in loss of enzymatic activity. Although the cytoplasmic cavity where the heme molecule is bound shows conservation in other MBOAT family members, the position of Cys-324 appears to be unique to HHAT. Could the HHAT-bound heme group have a regulatory role or is it present to block dimerization observed in other small molecule lipid MBOATs? A possibility could be that the heme-Cys324 binding site is a regulatory modification site that can compete with e.g. Cys-palmitoylation, as it has previously been shown that the C324A mutant affects the availability of *S*-palmitoylation sites within HHAT (Konitsiotis et al., 2015). Crosstalk between palmitoylation and heme attachment has not previously been reported, but would provide an appealing additional level of regulation as has been shown between palmitoylation and ubiquitination/phosphorylation in transmembrane proteins (Tian et al., 2008; Yount et al., 2012).

HHAT has been proposed to possess a transporter function for Palm-CoA (Asciolla and Resh, 2019). Our structural analysis reveals a distinct Palm-CoA binding site essential for acyl transfer that would not allow for a direct transport mechanism. However, we cannot rule out that when HHAT is in an apo state (without bound SHH or nanobody bound in the luminal cavity), a conformational change could occur that would allow the CoA palmitoyl chain to adopt an elongated conformation pointing towards the luminal side. Such different conformations of the acyl moiety were previously observed in the structures of DGAT (Fig. 2G), and further structural studies of HHAT apo state(s) will be required to determine the mechanism of a potential transport function.

We propose a model of HHAT function based on our structures (Fig. 5). In the absence of substrate, Trp335 blocks water permeation by obstructing the central solvent channel (Fig. S7). The lipid tail of Palm-CoA enters from the cytosolic face and Trp335 rotates into an ‘open’ conformation. This HHAT ‘priming’ allows the SHH N-terminus to bind inside the luminal cavity, and this binding is occluded by NB169 and 177 which compete with SHH for HHAT binding (Fig. 4E). We observe a larger density present in a luminal pocket of HHAT adjacent to the luminal cavity, consistent with a sterol moiety (Fig. S9). It is interesting to note, given the C-terminal cholesterylation unique to hedgehog homologues across all known proteins, that a cholesterol group could occupy the lipid density. Conserved Asp339 may form a charge interaction with the SHH N-terminal amine to orientate the residue for attack; this residue is Asn in PORCN and GOAT, which acylate internal residues THAT are more likely to be orientated by complimentary hydrogen bonding. It remains unclear whether *N*-palmitoylation is directly catalysed by HHAT, or whether it results from spontaneous rearrangement of a thiol-attached intermediate (Mann and Beachy, 2004). Palmitoylation will increase hydrophobicity of an already C-terminally cholesterylated SHH protein. Acylated SHH is therefore unlikely to exit HHAT directly into the cytosol and needs to detach via lateral exit into the membrane, and/or supported by secreted and membrane proteins (e.g. Scube2 and Dispatched). This requirement for assisted product release is supported by our previous observation that *N*-palmitoylated SHH or SHH N-terminal peptide are relatively potent inhibitors of HHAT under cell-free conditions (Lanyon-Hogg et al., 2019).

**Fig. 5.**
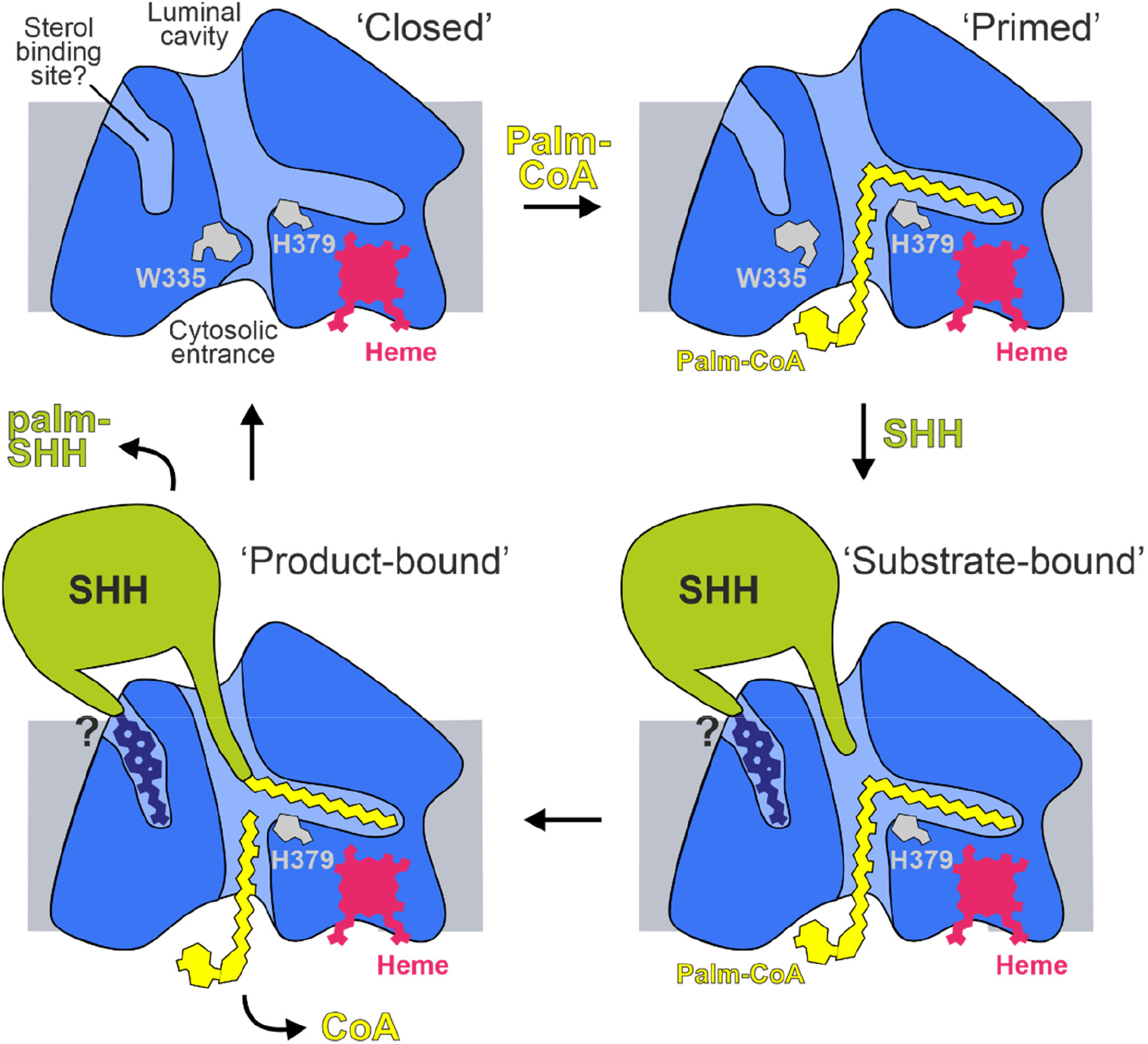
Model for HHAT-mediated palmitoylation of SHH. Without any substrate bound, HHAT adopts a ‘closed’ conformation with Trp335 blocking the Palm-CoA binding pocket (top left panel). Binding of Palm-CoA from the cytosolic side leads to a rearrangement of the active site involving residues His379 and Asp339 and primes HHAT for SHH binding (top right panel). The SHH N-terminus binds HHAT from the luminal cavity to access the active site (bottom right panel). SHH binding may be supported by binding of the C-terminal SHH-cholesterol moiety to a potential sterol binding pocket located adjacent to the luminal cavity. Once acyl transfer to SHH occurred, palmitoylated SHH detaches from HHAT via either lateral exit into the membrane and/or supporting secreted and membrane proteins (e.g. Scube2 and Dispatched) (bottom left panel).

Inhibiting HH signalling is a proven strategy in anti-cancer therapy. Most prior drug development has focused on developing inhibitors of the GPCR and HH signal transducer SMO, and SMO inhibitors are used in the clinic for the treatment of patients with basal cell carcinoma, medulloblastoma and acute myeloid leukemia (Wu et al., 2017). However, patients frequently develop mutations in SMO that cause drug resistance. Here, we provide a structural template for the development of HHAT inhibitors, a validated alternative strategy to inhibit SHH signalling. Moreover, our structure suggests new approaches to control lipid attachment to SHH, which could be used to engineer SHH morphogens with different specificities to manipulate stem cells and morphogen gradients.

### Future directions and limitations

The findings presented here offer crucial insights into the initial Palm-CoA loading step in the HHAT enzymatic cycle, and into inhibition of the reaction centre for future development of therapeutic agents. However, important questions about the HHAT mechanism will need to be addressed in future studies. A comprehensive understanding of the entire HHAT mechanism requires elucidation of intermediate conformations within the catalytic cycle via structural methods, alongside corresponding energetic profiles, such as can be obtained using quantum mechanical/molecular mechanics (QM/MM) methods. Discerning these intermediate structures will require astute employment of advancing cryo-EM methods, such as the use of conformation-specific nanobodies/megabodies to trap scarce protein subpopulations and/or the use of small-molecule mimetics of reaction intermediates. QM/MM calculations are currently limited by the size of the simulated system, making application to enzymes acting on complex lipidic substrates computationally challenging (Medina et al., 2018).

Determination of the structure of HHAT bound to its substrate SHH may also assist interpretation of densities protruding into HHAT, including whether the proposed sterol binding pocket (Fig. S9) could be occupied by the C-terminal cholesteryl moiety of SHH (Fig. 5). One prevailing question relates to whether palmitoylated-SHH exits HHAT either by moving laterally into the surrounding membrane or via ‘upwards’ dissociation into the luminal solvent. A first glimpse of a putative product exit process is offered by simulations where we observe spontaneous and stable binding of a lipid molecule between helices α5 and α9 of HHAT, whereby one lipid acyl-tail mimics the proposed position of palmitoylated-SHH in the reaction centre (Fig. S10**, Movie S1**). Binding of the product mimetic lipid-tail is accompanied by outward movement of α5 which correlates with inwards swivel of Tyr377 to open the ‘luminal gate’ (Fig. S10A-G). Removal of lipid from the gate region or via pulling the lipid through the gate enabled relaxation of the α5-α9 distance, accompanied by reset and outward swivel of the Tyr377 sidechain (Fig. S10F-H). These data offer tantalising clues as to the potential mechanism of palmitoyl-SHH unloading (Fig. S10I) into the membrane which will be investigated in future experiments. Product exit studies will also need to account for whether the amine group of SHH-Cys24 is directly palmitoylated, or whether the reaction proceeds via initial *S*-palmitoylation followed by *S*,*N* acyl rearrangement. Thus, our current studies point the way from structure determination to an atomistic mechanistic understanding of HHAT function in a complex cell membrane environment. Finally, future studies should translate these mechanistic insights into a cellular context, and the impact on SHH maturation and signalling.

Whilst our work was in preparation for submission, a structure of HHAT was disclosed by Jiang *et al*. (Jiang et al., 2021); we discuss the relationship to the structures reported here, and highlight similarities and distinctive differences between them in Fig. S11 and Supplementary Results and Discussion.

## ACKNOWLEDGEMENTS

We thank the staff at the DLS UK National Electron Bio-Imaging Centre (eBIC) for assistance with cryo-EM data collection and S. Scott and D. Staunton for technical support. We thank Eva Beke for technical assistance and Megabody constructions. We also thank C. D. Johnston and J. Newington (Imperial College London) for analysis of nanobodies. Work was supported by Cancer Research UK (C20724/A14414 and C20724/A26752 to C.S.), European Research Council (647278 to C.S.), BBSRC (BB/T01508X/1 to E.W.T. and C.S, BB/R00126X/1 to M.S.P.S.), the European Union Horizon 2020 programme (Marie Skłodowska-Curie Individual Fellowship 101026939 to S. A. and E.W.T.) and Wellcome (Technology Development Grant 208361/Z/17/Z to M.S.P.S., DPhil studentships 102749/Z/13/Z to L.S., and 102164/Z/13/Z to C.E.C. and T.B.A.). We acknowledge the support and the use of resources of Instruct-ERIC (PID3603 & PID10022), part of the European Strategy Forum on Research Infrastructures (ESFRI), and the Research Foundation - Flanders (FWO). This project made use of time on ARCHER via the HECBioSim, supported by EPSRC (EP/L000253/1). Electron microscopy provision was provided through eBIC (proposal EM20223) and the OPIC Electron Microscopy Facility (funded by Wellcome JIF (060208/Z/00/Z) and equipment (093305/Z/10/Z) grants). The computational aspects of this research were funded from the NIHR Oxford BRC with additional support from the Wellcome Trust Core Award Grant Number 203141/Z/16/Z. The views expressed are those of the author(s) and not necessarily those of the NHS, the NIHR or the Department of Health. Further support from the Wellcome Core Award 090532/Z/09/Z and BHF (RE/18/3/34214).is acknowledged.

## AUTHOR CONTRIBUTIONS

C.S. and E.W.T. designed the project. C.E.C, P.K., L.S. and E.F.X.B. expressed and purified the proteins for cryo-EM analysis, biophysical experiments, enzymatic assays and nanobody immunisation. C.E.C. carried out BLI, UV/VIS spectroscopy and thermostability analyses, P.K. carried out cloning and production of the HHAT mutants with assistance from C.E.C. S. A. A. performed enzymatic assays. R.A.S. performed immunodetection of HHAT-nanobody interactions. L.C. and C.E.C. collected and processed electron microscopy data, L.C., C.S. and C.E.C. built and refined models. T.B.A. and M.S.P.S performed MD simulations and analysis. E.P. and J.S. designed and generated nanobodies and megabodies, and C.E.C. performed nanobody screening, expression and purification. A.I.M. and T.L.-H.analysed the data and contributed to writing the manuscript. C.S. and E.W.T. supervised the studies, analysed data, and prepared the manuscript with input from all authors.

## SUPPLEMENTARY FIGURES S1-S11 AND TABLE S1

**Fig. S1.**
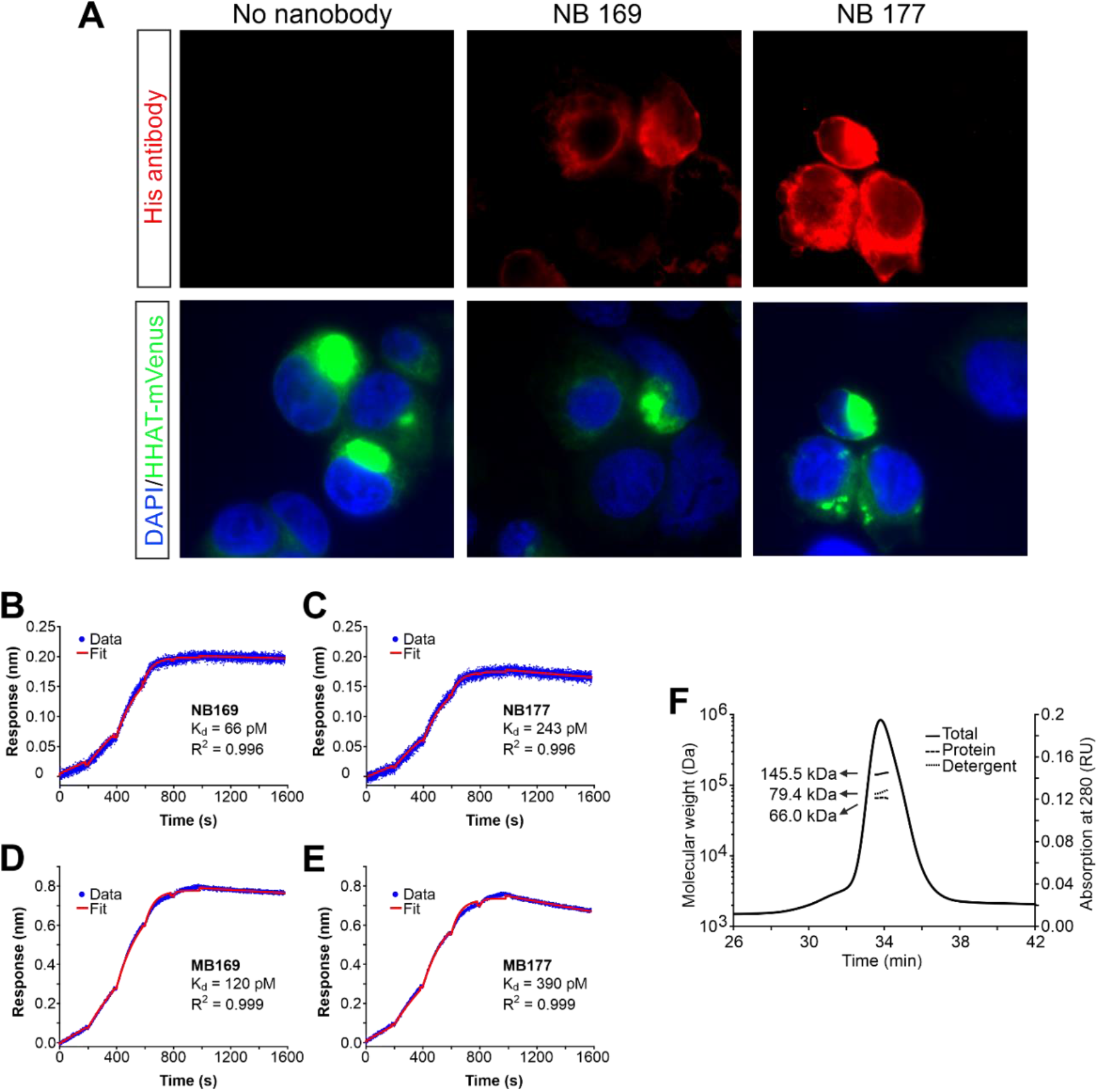
Functional characterisation of HHAT-nanobody and megabody interactions. Related to Figure 1. **(A)** Immunofluorescence staining of mVenus-tagged full-length human HHAT stably expressed in COS-7 cells (green). The figure shows representative images of HHAT-mVenus expressing cells in green, incubated without or with either NB169 or NB177. Nanobodies were visualised by staining with primary and secondary antibodies and are shown in red. These analyses showed co-localisation between mVenus-tagged HHAT and NB staining, indicating that both nanobodies readily bind to HHAT embedded in the cellular membrane. (**B-E),** Biolayer interferometry (BLI) of HHAT nanobody and megabody interactions. Single cycle kinetic measurements for the HHAT-NB169 (**B**), HHAT-NB177 (**C**), HHAT-MB169 (**D**) and HHAT-MB177 (**E**) interactions are shown. K_d_ and R^2^ are indicated for each measurement **(F)** SEC-MALS analysis of OGNG-solubilised human HHAT. Molecular weights (MW, black lines) and 280 nm absorption (grey line) plotted against elution time. For clarity, graphs of MW are shown only around main absorption peak. Theoretical MW of HHAT based on sequence and including heme-B and 6 palmitoylation sites is 60.3 kDa. This analysis suggests that human HHAT is a monomer under our purification conditions.

**Fig. S2.**
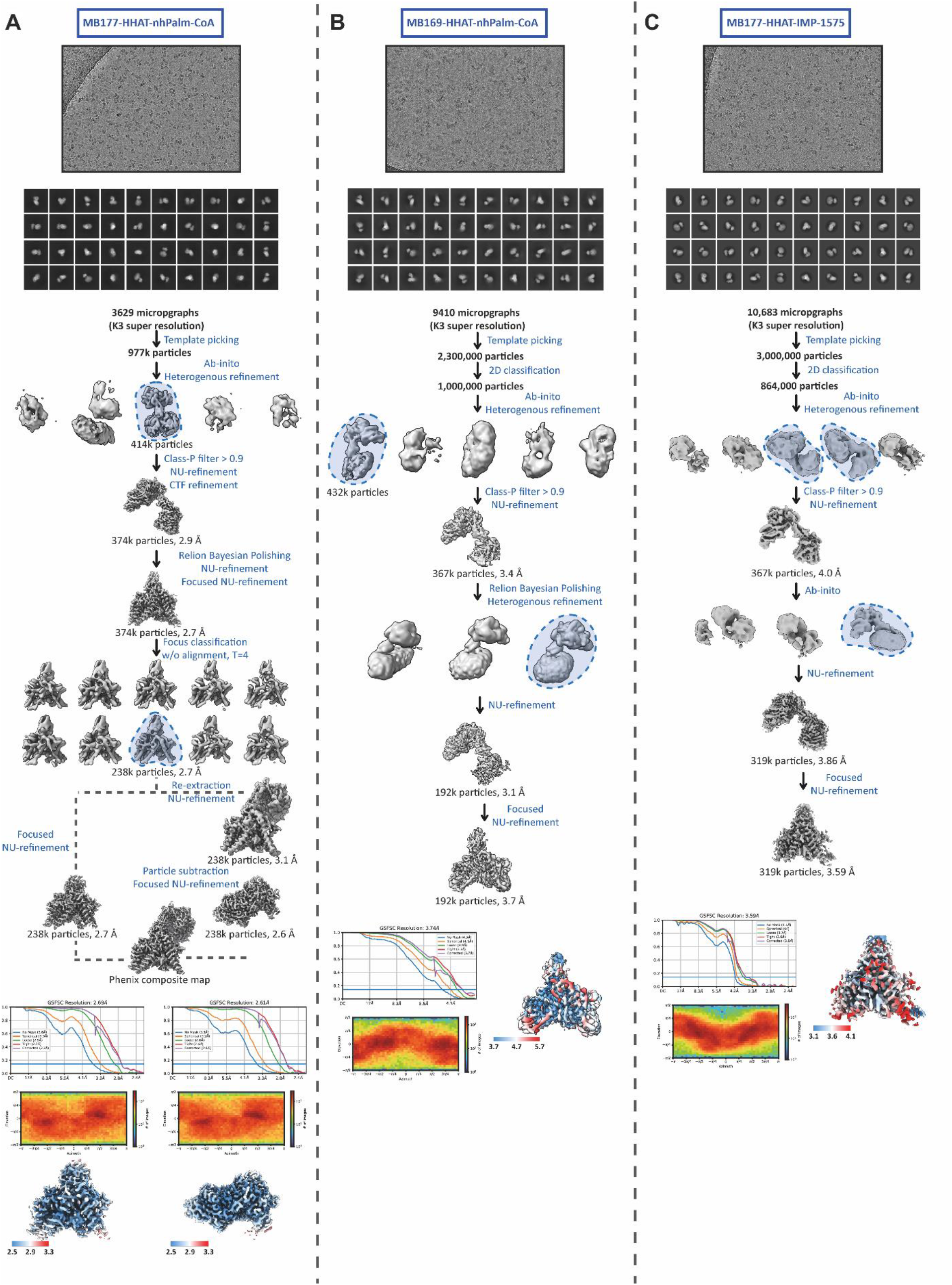
Cryo-EM data collection, processing and analysis scheme. Related to Figures 1-4. Flowchart for the processing and the classification of the HHAT-MB270-nhPalm-CoA (**A**), HHAT-MB169-nhPalm-CoA (**B**) and HHAT-MB270-IMP-1575 (**C**) complexes. For panel, the top figure is a micrograph followed by 2D classes, the processing flowchart with the FSC plot and final angular distribution as well as the local resolution estimation.

**Fig. S3.**
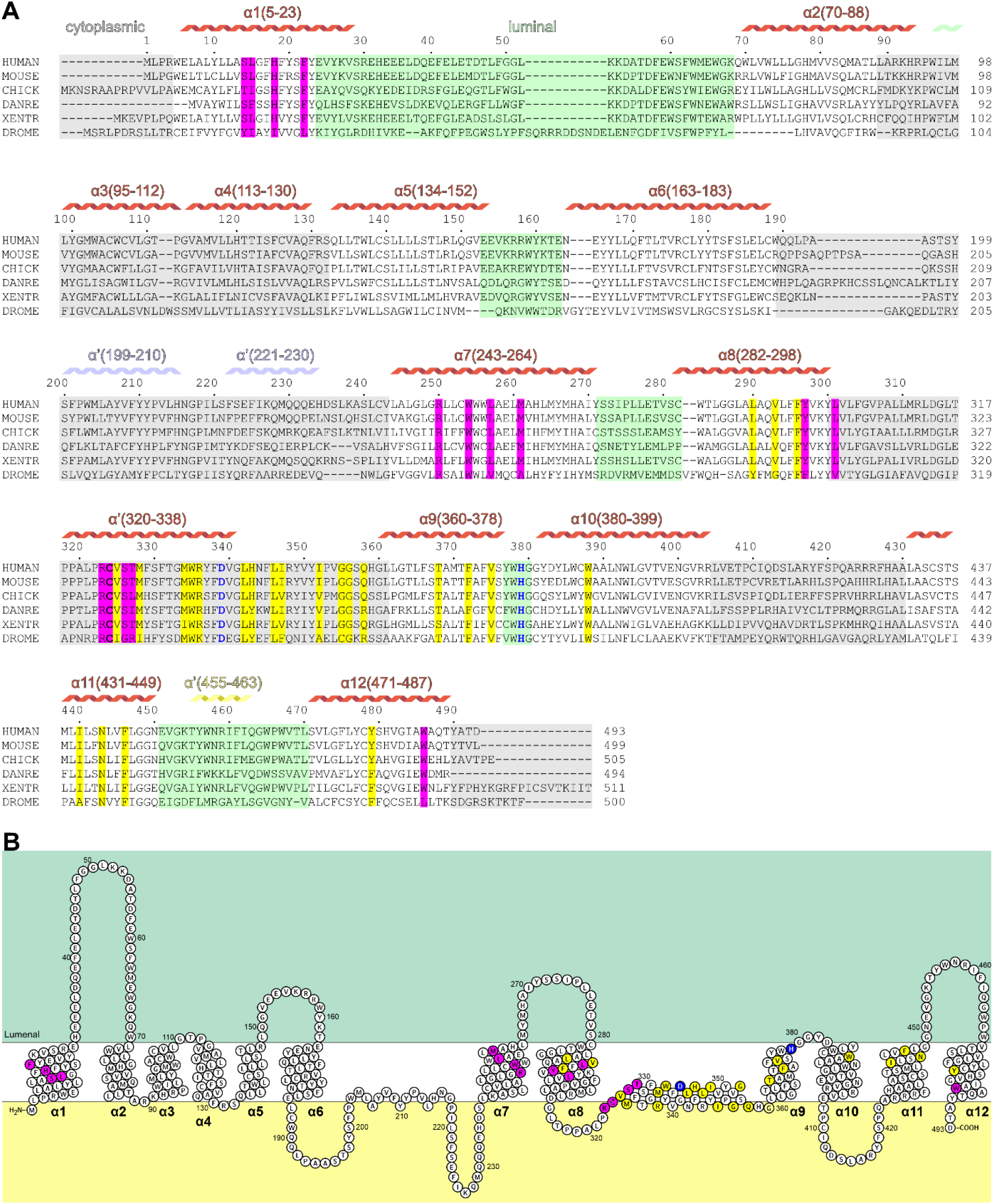
Sequence alignment and topology of HHAT. Related to Figures 1-3. **(A)** Sequence alignment of selected HHAT family members. Secondary structure assignment is displayed above the alignment. Grey boxes correspond to cytosolic and green boxes to ER luminal loop regions. Residues interacting with heme-B are in magenta and with nhPalm-CoA in yellow. Active site residues Asp339 and His379 are highlighted in blue. **(B)** Topology diagram of human HHAT adapted from the Protter server (wlab.ethz.ch/protter/start/) and colour-coded as in **A**. UniProt Ids are as follows: Q5VTY9 (human), Q8BMT9 (mouse), F1NHW1 (chick), F8W2J4 (zebrafish, DANRE), F6YU00 (frog, XENTR), Q9VZU2 (fly, DROME).

**Fig. S4.**
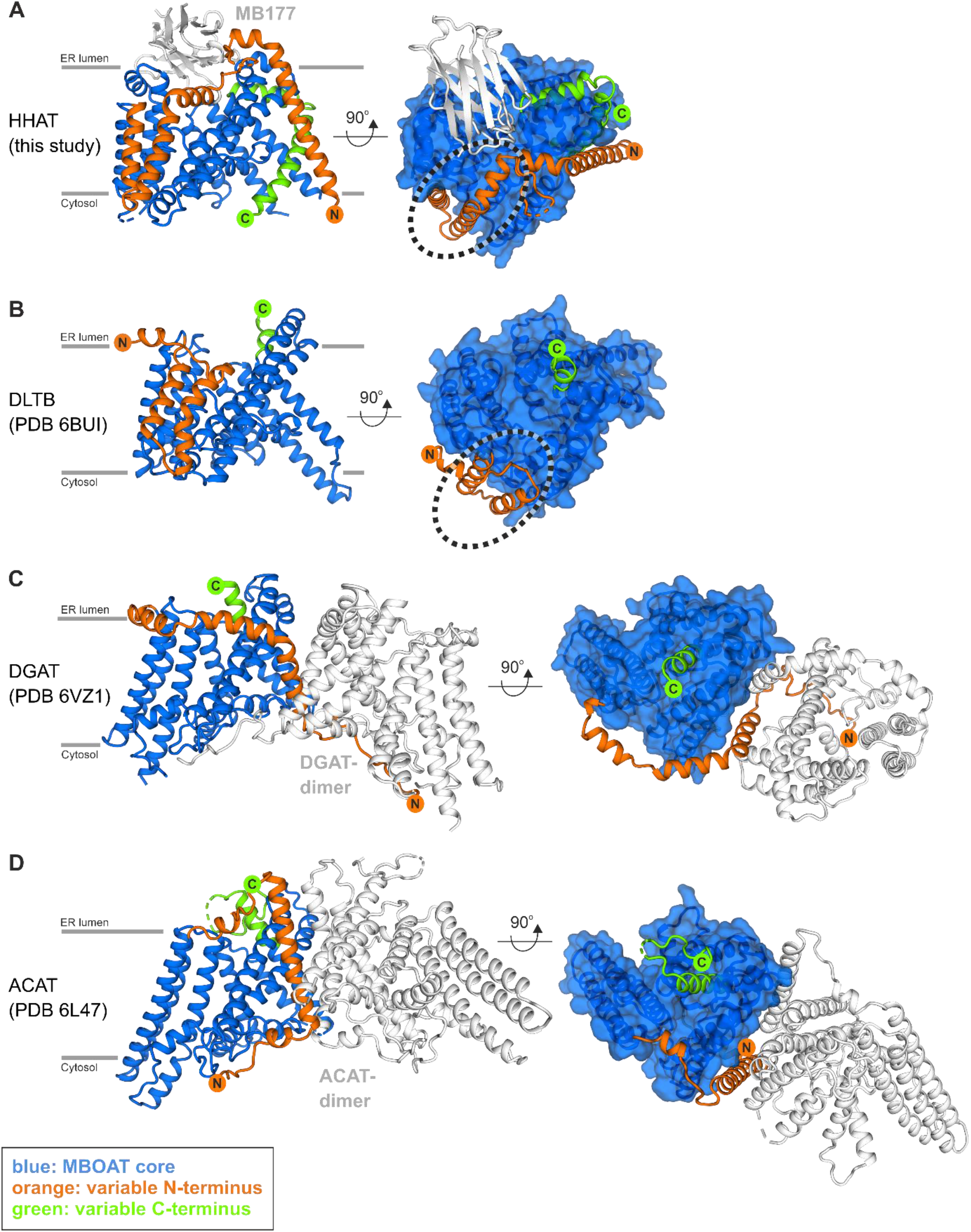
Structural comparison of MBOAT superfamily members. Related to Figure 1. Cartoon representations of HHAT (**A**), DLTB (**B**, PDB Id. 6BUI), DGAT (**C**, PDB Id. 6VZ1) and ACAT (**D**, PDB Id. 6L47). DLTB, DGAT and ACAT were superimposed onto HHAT. The conserved MBOAT core is coloured in blue, and the variable N- and C-termini in orange and green, respectively. The dotted circles in the right panels of **A** and **B** indicate conserved transmembrane helices in HHAT and DLTB, that are rearranged in DGAT and ACAT to participate in dimer formation.

**Fig. S5.**
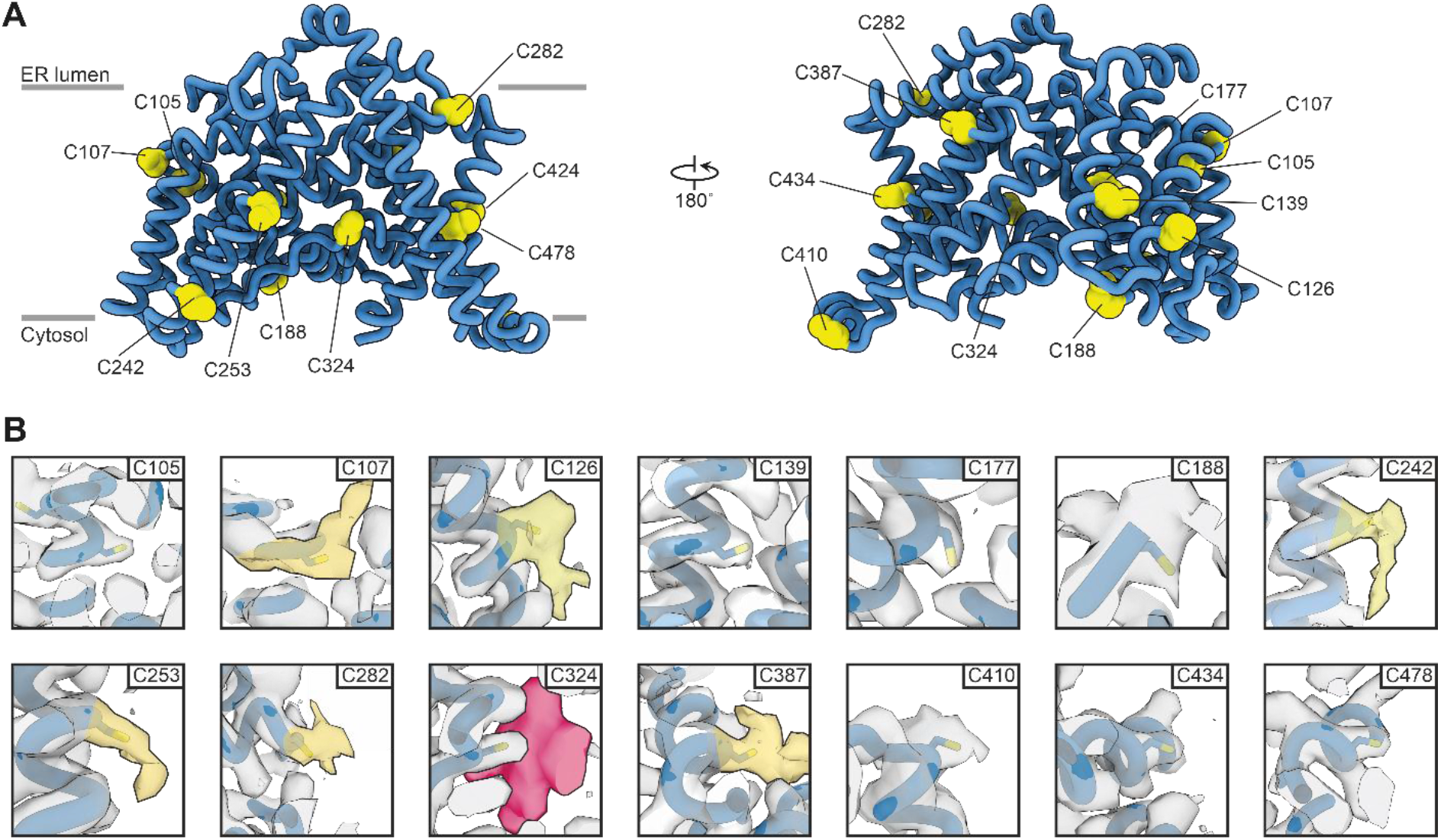
Analysis of potential HHAT cysteine palmitoylation sites. Related to Figure 1. **(A)** Cartoon representation of HHAT with all cysteine residues labelled and highlighted as yellow spheres. **(B)** Close-up views of all cysteine residues. The high resolution cryo-EM map is shown in grey. Observed additional density is highlighted in yellow. The density around side chain C324 that we identified as a heme-B group is depicted in red. We note that Cys-188 is the terminal residue of a disordered, cytoplasmic loop and thus identification of additional map features that could correspond to palmitoylation are difficult to access.

**Fig. S6.**
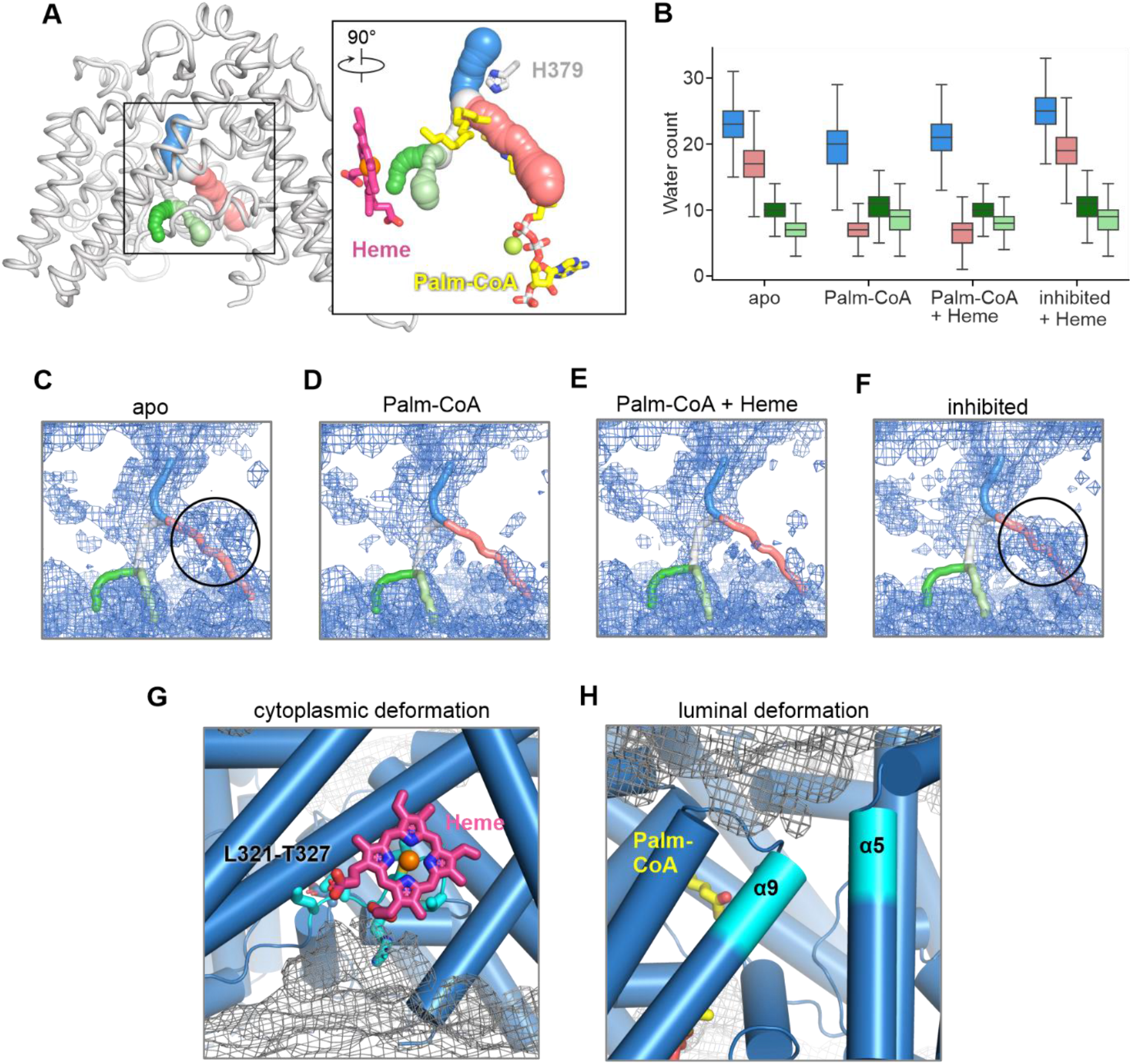
Water and membrane densities in HHAT simulations. Related to Fig. 2. **(A)** Tunnels through the HHAT core as calculated using Caver. Tunnels are displayed as spheres. The inset indicates the position of tunnels with respect to the bound Heme (pink), Palm-CoA (yellow) and His379 (grey) which are displayed as sticks. The Fe and Mg ions are shown as spheres in orange and green, respectively**. (B)** Quantification of the number of water within 4 Å of the centroid coordinates through the tunnels indicated in **A** across the final 50 ns of atomistic MD simulations of HHAT in the apo, Palm-CoA, Palm-CoA plus HEME and inhibited/Heme bound states. Boxplots are coloured according to the tunnel colours in **A**. **(C-F)** Time averaged solvent density within the HHAT core across atomistic simulations of HHAT in the apo (**C**), Palm-CoA bound (**D**), Palm-CoA plus Heme (**E**) and inhibited/Heme bound (**F**) conformations. Trajectories were combined and fitted before the density calculations which were performed using MDAnalysis. The centroid through the tunnels are shown as sticks and the circles indicates solvent within the Palm-CoA binding pocket. **(G-H)** Closeup of the regions of membrane deformation in the Heme binding cavity and around α5/α9 (the luminal gate). Cyan sticks depict the location of a loop formed by Leu321-Thr327 which includes the Heme coordinating Cys324. Membrane density (grey isomesh) was calculated in CG simulations of HHAT and overlayed with the atomistic structure for reference.

**Fig S7.**
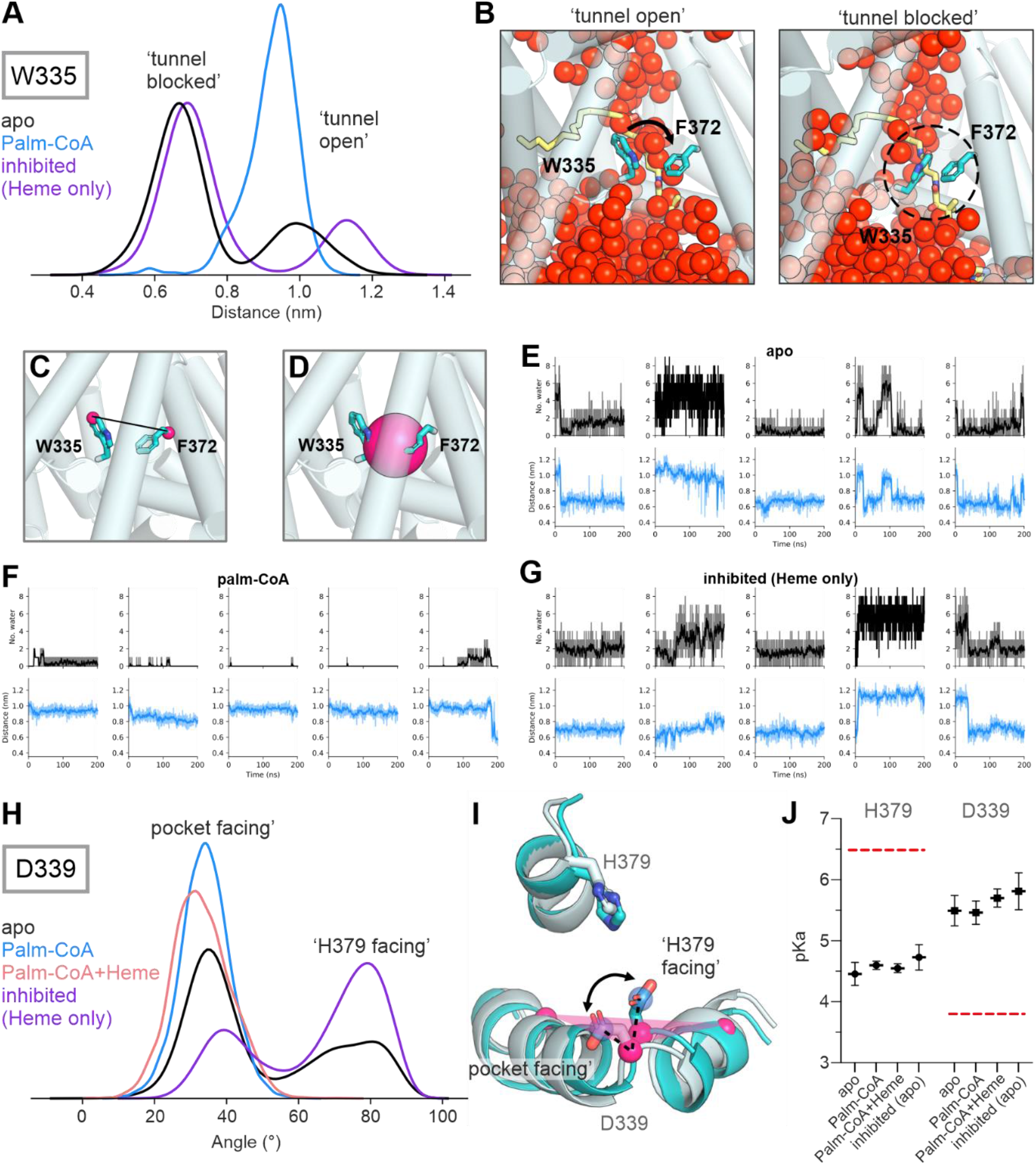
HHAT core dynamics. a-g, Trp335 dynamics and water permeation. Related to Fig. 3. **(A)** Distance between the Cη2 atom of Trp335 and the Cα atom of Phe372 across simulations of HHAT in the apo, Palm-CoA bound and inhibited/Heme only states. **(B)** Snapshots showing the conformation of Trp335 at the start (‘tunnel open’) and end (‘tunnel blocked’) of a simulation of HHAT apo. Trp335 and Phe372 are shown as cyan sticks and coloured by atom, HHAT is shown as transparent cylinders and water atoms are shown as red spheres. Snapshots are overlaid with the position of Palm-CoA (yellow sticks) to indicate the relative position of the binding pocket. **(C)** Close-up view of the distance between the Cη2 bead of Trp335 and Cα bead of Phe372 (pink spheres). **(D)** Sphere with a radius of 4Å, positioned at the centre of geometry of Trp335/Phe372 Cα atoms. Trp335/Phe372 are shown as sticks and HHAT is shown as transparent cylinders. **(E-G)** Number of waters within the sphere (black) and Trp335/Phe372 distance (blue) vs time in simulations of HHAT in the apo (**E**), Palm-CoA bound (**F**) and inhibited/Heme bound (**G**) conformations. MDAnalysis was used to calculate the position of the sphere at each time point in the trajectory such that the spheres position varied according to the centre of geometry of the Trp335/Phe372 Cα’s. The rolling mean is shown as an opaque line against the raw data (transparent) to reduce noise resulting from Brownian fluctuations. **(H-J),** Dynamics of the HHAT reaction centre. **(H)** Angle of the Asp339 sidechain across atomistic MD simulations of HHAT in an ER mimetic bilayer. Simulations (5x 200 ns of each) were initiated from either the HHAT-Palm-CoA-Heme structure in apo (black), Palm-CoA bound only (blue) or Palm-CoA bound plus Heme (salmon red) states, or from the inhibited HHAT structure with the inhibitor removed such that only the Heme was bound (purple). The Asp339 sidechain angle was defined as the angle between a vector formed by the Cα and Cγ atoms of Asp339 with respect to a plane formed by the Cα atoms of Met334, Asp339 and Leu346. **(I)** Overlay of the start (light cyan) and end (cyan) snapshots from a simulation of HHAT in the apo state, indicating the change in position of Asp339 between the ‘pocket facing’ and ‘His379 facing’ conformations. Atoms used to define the vector (blue) and plane (pink) are shown as spheres and the vector is indicated by a dashed black line. **(J)** The pKa of Asp339 and His379 across simulations of HHAT in the apo, Palm-CoA, Palm-CoA plus Heme and inhibited/Heme only simulations as calculated using pKa-traj. Red dashed lines indicate standard pKa values of histidine or aspartate residues. pKa values are shown as the mean value and error bars indicate the standard error of the mean between replicates.

**Fig. S8.**
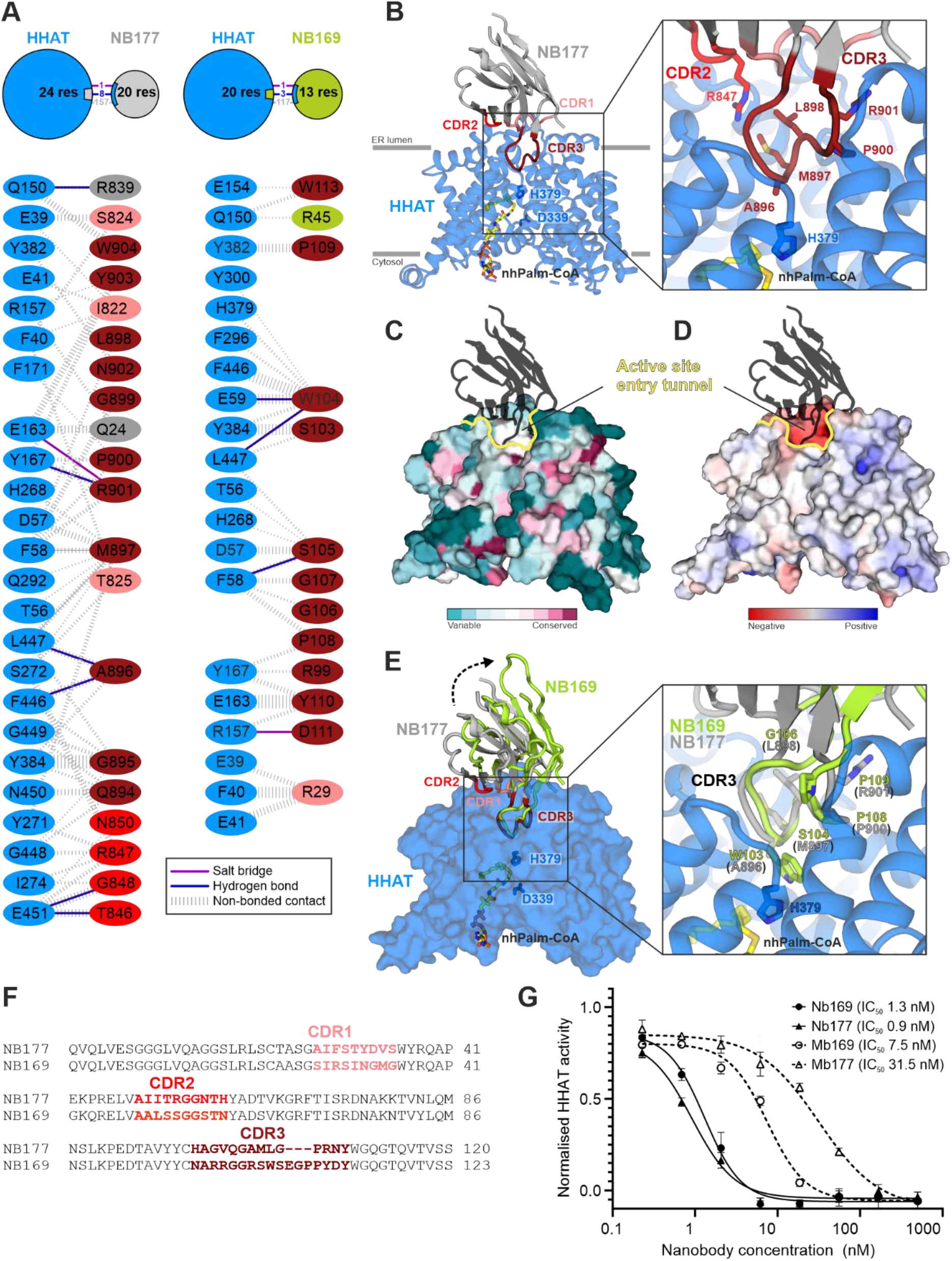
Structural analysis of the HHAT-MB177 complex. Related to Figure 4. **(A)** Schematics of the HHAT-NB177 interactions observed in the high resolution cryo-EM structure (adapted from PDBSUM (REF)). HHAT is coloured in blue, NB177 in grey with the complementary determining regions (CDRs) coloured in salmon (CDR1), red (CDR2) and dark red (CDR3). **(B)** Cartoon representation of the HHAT-NB177 complex, colour-coded as in **A**. Active site HHAT-residues His379 is highlighted in stick representation, The box shows a close-up of the active site entry tunnel that is mainly blocked by NB177-CDR3, that protrudes deeply into HHAT. **(C-D)** Surface representation of HHAT with NB177 shown in black ribbon. Sequence conservation (**C,** calculated with Consurf) and electrostatic potential (**D**, from ± 8kT/eV) are mapped onto the solvent accessible HHAT surface. The active site entry tunnel is outlined in yellow. **(E)** Structure of the HHAT-NB169 complex. HHAT is shown as blue surface and NB169 in green cartoon representation. NB177 is shown superimposed and coloured as in a and b (using HHAT as template). Orientation is as in b. A close-up of CDR3 is shown in the box, revealing the close structural similarity of the Cα backbone. **(F)** Sequence alignment of nanobodies NB177 and NB169 with the complementary determining regions (CDRs) coloured as in a. Overall sequence identity is 77.4 %, with only 27.8% identity in the CDRs (CDR1: 22.2 %, CDR2: 40.0 %, CDR3: 23.5 %). **(G)** Acyl-cLIP assay to test SHH palmitoylation. Dose response curves of nanobodies and megabodies inhibiting SHH palmitoylation by HHAT. The IC50 of the nanobodies is below the detection limit of the assay ([HHAT] ≈ 10 nM). N = 1 (triplicates), mean ± SEM.

**Fig. S9.**
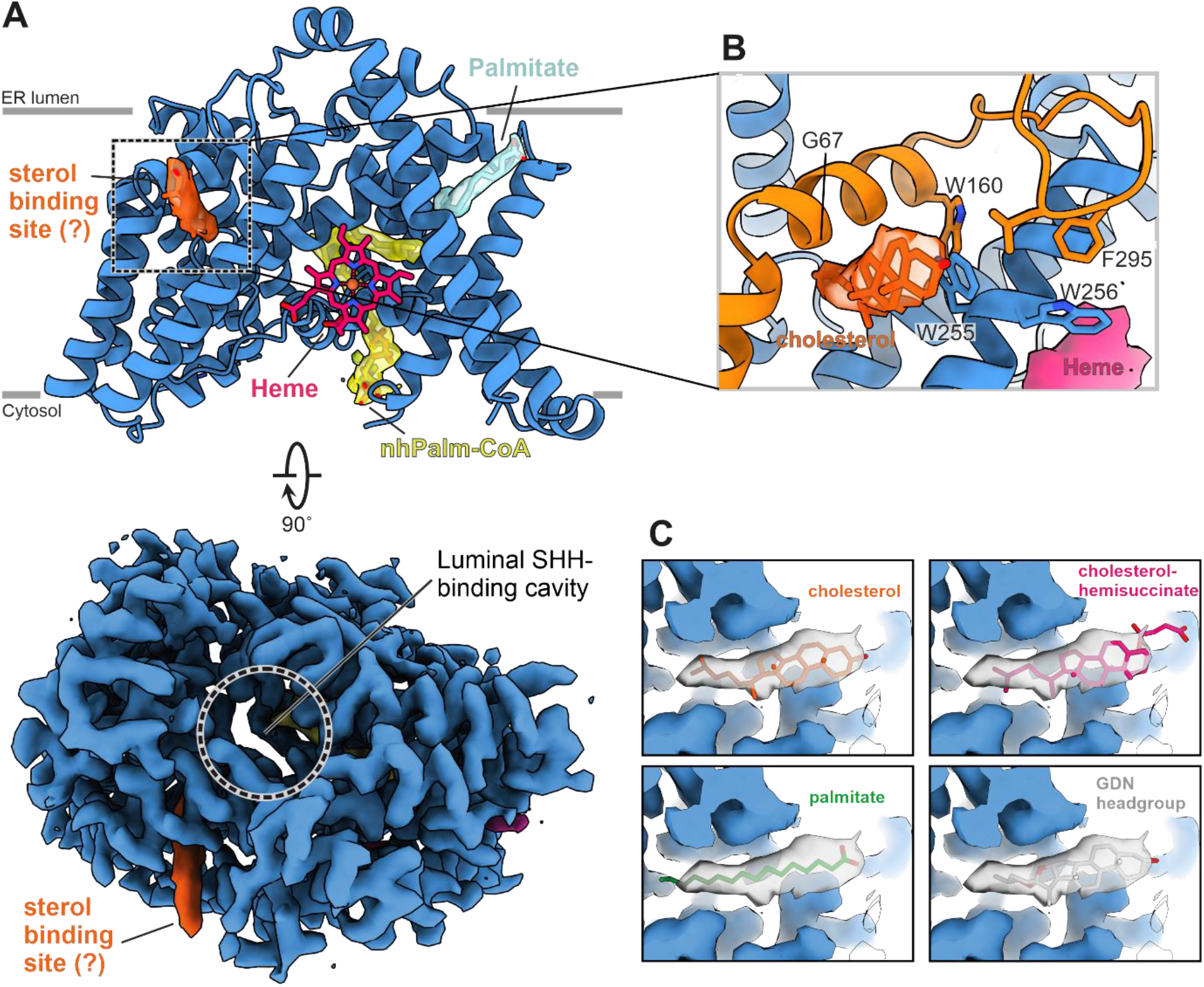
A potential sterol binding site adjacent to the luminal SHH binding cavity. Related to Figure 4. **(A)**, Cartoon presentation of HHAT with selected lipids and substrates highlighted and depicted and drawn with the final cryo-EM map. View and colour coding is as in Fig. 2. **(B)** Close-up view of the potential sterol-binding site. The final cryo-EM (orange) is suggestive of a cholesterol or similar sterol molecule. **(C)** Comparison of the different lipids fitted into the sterol map shown in **B**.

**Fig. S10.**
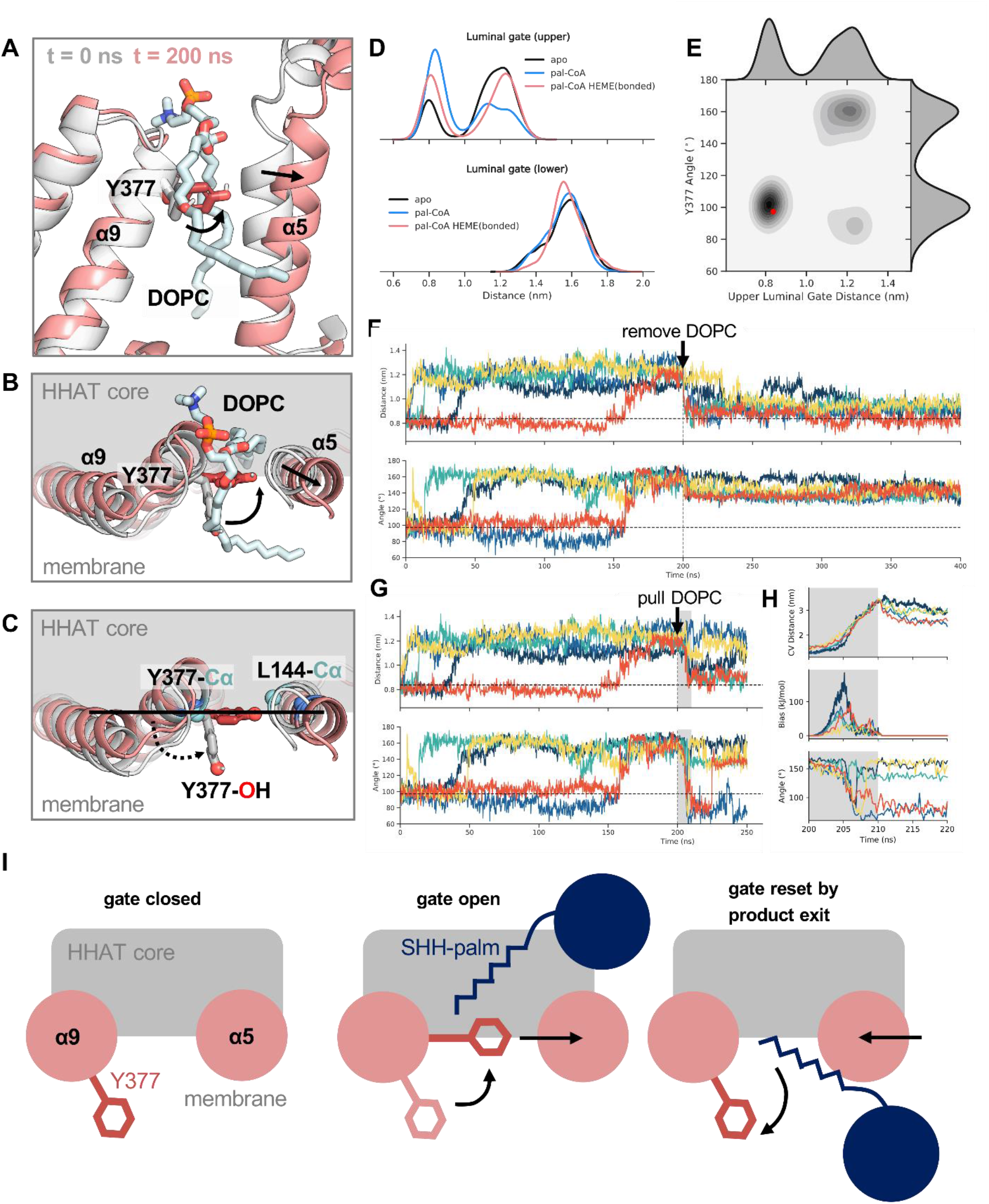
A luminal exit gate for palmitoyl-SHH. **(A)** Snapshots from the start (t = 0 ns, grey) and end (t = 200 ns, red) of an atomistic simulation of HHAT, with a DOPC lipid tail (light blue sticks) located in the luminal gate mimicking palmitoylated-SHH product. Arrows indicate outward movement of α5 and inward swivel of the Tyr377 sidechain (shown in stick representation). Atomistic simulations were initiated from a coarse-grain simulation snapshot where DOPC spontaneously and stably bound within the luminal gate (**Movie S1**). Backmapping to atomistic resolution was achieved using CG2AT (https://github.com/owenvickery/cg2at). **(B)** Top view of the luminal surface showing expansion of the luminal gate and change to the sidechain angle of Tyr377 when one acyl-tail of the DOPC lipid is positioned within the reaction centre. **(C)** Definition of the upper luminal gate distance and Tyr377 sidechain angle distributions analysed in **D-H**. The upper luminal gate distance was defined as the distance between Cα atoms of Try377/Leu144 (cyan spheres). The Tyr377 sidechain angle was defined as the angle between a vector formed by the Cα and Oh (hydroxyl) atoms of Tyr377 and a vector formed by the Cα atoms of Tyr377/Leu144 (shown as spheres). **(D)** Distribution of distances at the top/bottom of α5/α9 across 5 x 200 ns atomistic simulations of apo HHAT (black), HHAT with palm-CoA bound (blue) and HHAT with heme and palm-CoA bound (red). All simulations were initiated with DOPC bound in the luminal gate indicating consistent opening of the top of the luminal exit gate due to outward movement of α5. **(E)** 2D correlation plot of the upper luminal gate distance vs the Tyr377 sidechain angle (as defined in **C**) across 5 x 200 ns simulations of apo HHAT. The red dot indicates the distance and angle at t = 0 ns. **(F)** Upper luminal gate distance and Tyr377 sidechain angle with time indicating correlated motions across the first 5 x 200 ns of apo HHAT simulations. At 200 ns the bound DOPC was removed from the luminal gate and a further 200 ns of atomistic simulation performed for each replicate to allow relaxation of the gate. (**G**) As in F but at 200 ns the DOPC was pulled out of the luminal gate into the membrane over a period of 10 ns (grey box) before a further 40 ns of unbiased atomistic simulation was performed for each replicate. The collective variable (CV) used during the steered MD simulations was defined as the distance between the centre of mass of the DOPC lipid and the Cα atom of Arg176. A force constant of k = 1000 kJ mol^-1^ nm^-2^ was gradually increased over the 10 ns steered MD simulation using PLUMED (https://www.plumed.org). (**H**) Zoomed in view of the steered MD simulation (grey box) from **G** indicating exit of DOPC through the gate as shown by the CV distance (top) as the work bias factor is increased (middle) and the Tyr377 sidechain angle swings outward (bottom) to permit DOPC exit. (**I**) Model of the proposed luminal exit gate mechanism for palmitoylated-SHH. In the absence of palmitoylated-SHH product the luminal gate formed by α5/α9 is closed. When palmitoylated-SHH is located within HHAT, the upper region of α5 tils outwards allowing the sidechain of Tyr377 to swing inwards and plug the gate. As palmitoylated-SHH moves laterally into the membrane the Tyr377 sidechain is pushed outwards, enabling inward movement of α5 to reset the luminal gate.

**Figure S11.**
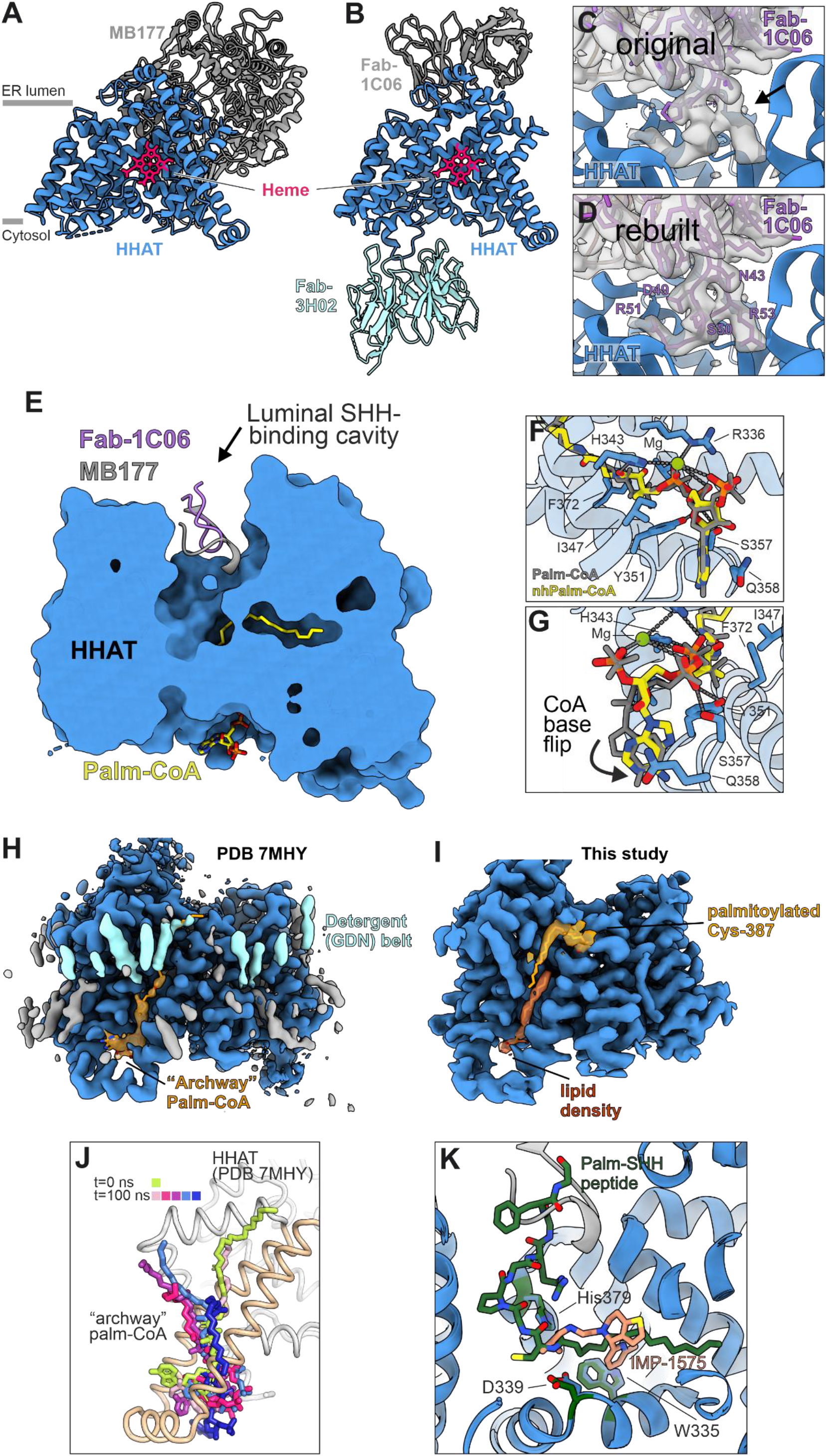
Structural comparison to the HHAT-Fab1C06-Fab3H02 complex from Jiang et al. **(A, B)** Cartoon representations of the HHAT-MB177 complex (this study) and the HHAT-Fab1C06-Fab3H02 complex (Jiang *et al*). Structures were superimposed using HHAT as template. **(C, D)** Close-up view on the Fab1C06-HHAT interface showing the 2.7 Å map (EMD-23836), and original (**C**, 7MHY) and rebuilt (**D**) PDB file. **(E)** HHAT solvent-accessible surface with the HHAT interacting regions of MB177 (this study) and Fab1C06 (Jiang et al) shown in cartoon representation and nhPalm-CoA in sticks. **(F, G)** Two close-up views of the Palm-CoA base region. nhPalm-CoA (this study) is shown in atomic colouring (yellow: carbon, red: oxygen, blue:nitrogen, orange: phosphor), the Palm-CoA from Jiang *et al*. in grey. The movement of the base region is indicated with an arrow. (**H, I**) Map comparison of the “archway” Palm-CoA region between Jiang *et al*. (**H**, EMD-23836) and this study (**I**). The detergent (GDN) belt is coloured in cyan, the “archway Palm-CoA” in light orange and the palmitoylated Cys-387 in dark orange. **(J)** Comparison of the position of the “archway” Palm-CoA as modelled by Jiang *et al*. (t = 0 ns, lime) or after 5 x 100 ns atomistic simulations (t = 100 ns, blues/purples). Simulations were performed using the HHAT structure from Jiang *et al*. bound to heme and both Palm-CoA molecules (PDB ID: 7MHY). Palm-CoA is shown as sticks bound within the proposed “archway” (light brown) shown in cartoon representation. **(K)** Close-up view of the reaction centre. Superposition of the HHAT-Palm-SHH peptide (green, Jiang *et al*, PDB 7MHZ) and HHAT-IMP-1575 (blue/salmon, this study) complexes. Both IMP-1575 and the Palm-SHH peptide induce a similar rearrangement of HHAT active site residues His379 and Asp339, and “gate keeper” residue Trp335.

**Table S1.** Cryo-EM data collection, refinement and validation statistics. Related to Figures 1-4.

|  | HHAT-MB177-nhpalm-CoA<br>EMD-XXXX<br>XXXX | HHAT-MB169-nhpalm-CoA<br>EMD-YYYY<br>YYYY | HHAT-MB177-IMP<br>EMD-ZZZZ<br>ZZZZ |
| --- | --- | --- | --- |
| **Data collection** |  |  |  |
| Microscope | Titan Krios IV (eBIC) | Titan Krios IV (eBIC) | Titan Krios I (eBIC) |
| Voltage (kV) | 300 | 300 | 300 |
| Detector | Gatan K3 with EF | Gatan K3 with EF | Gatan K3 with EF |
| Recording mode | Super Resolution | Super Resolution | Super Resolution |
| Magnification | 105,000 | 105,000 | 105,000 |
| Movie/micrograph pixel size (Å) | 0.4145 | 0.4145 | 0.4155 |
| Dose rate (e-/px/sec) | 15.1 | 13.9 2.3 | 1 |
| Number of frames per movie | 50 | 50 | 50 |
| Movie exposure time (s) | 2.5 | 3 | 3 |
| Total dose (e-/Å<sup>2</sup>) | 54.76 | 60.6 | 53.48 |
| Defocus range (um) | 1.0 to 2.5 | 1.0 to 2.5 | 1.0 to 2.5 |
| Volta Phase Plate | no | no | no |
| **EM data processing** |  |  |  |
| Number of movies/micrographs | 3637 | 9410 | 10,683 |
| Box size (px) | 400 | 400 | 400 |
| Particle number (total) | 977K | 2,300,000 | 3,000,000 |
| Particle number (post 2D) | / | 1,003,000 | 864K |
| Particle number (post 3D) | 414K | 367K | 319K |
| Particle number (used in final map) | 238K | 192k | 234K |
| Symmetry | C1 | C1 | C1 |
| Map resolution (FSC 0.143) | 2.7 | 3.74 | 3.65 |
| Local resolution range (FSC 0.5) | 2.45-6 | 3.71-10 | 3.07-6 |
| Map sharpening B-factor (Å<sup>2</sup>) | -56 | -125 | -152 |
| **Model Building and Validation** |  |  |  |
| Initial model used | De novo | XXXX | XXXX |
| Model composition |  |  |  |
| Non-hydrogen protein atoms | 21974 | 10178 | 9887 |
| Protein residues | 1357 | 606 | 597 |
| Ligands |  |  |  |
| nhpalm-CoA | 1 | 1 | / |
| IMP | / | / | 1 |
| Heme-b | 1 | 1 | 1 |
| CLR | 1 | 1 | 1 |
| Palm | 4 | 1 | 1 |
| Mg | 1 | 0 | 0 |
| RMSD from ideal |  |  |  |
| Bond length (Å) | 0.003 | 0.004 | 0.003 |
| Bond angles (°) | 0.558 | 0.663 | 0.593 |
| Validation |  |  |  |
| Molprobity score | 1.39 | 1.92 | 1.51 |
| Clashscore | 4.42 | 11.47 | 6.39 |
| Rotamers outliers (%) | 0.61 | 0 | 0 |
| FSC (0.5) model-vs-map | 2.7 | 4 | 3.7 |
| CC model-vs-map (masked) | 0.8 | 0.65 | 0.71 |
| Ramachandran plot |  |  |  |
| Favored (%) | 97.03 | 95 | 97.11 |
| Allowed (%) | 2.97 | 5 | 2.89 |
| Outliers (%) | 0 | 0 | 0 |

## SUPPLEMENTAL RESULTS AND DISCUSSION

### Comparison of our structure-function analysis to Jiang *et al*

Whilst preparing the final version of our manuscript, a structure of HHAT was disclosed by Jiang et al.(Jiang et al., 2021) (publication date 11 June 2021). We here present a comparative analysis of this structure with our own, and point out similarities and discrepancies. The cryo-EM maps obtained in both studies are of similar quality with an almost identical overall fold of HHAT (rmsd: 0.62 Å for 468 equivalent Cα positions) and a conserved binding mode for the previously uncharacterized heme-B coordinated by Cys324 (**Fig. S11A and B**). Both structures were obtained in complex with either an inhibitory megabody (MB177, Coupland et al.) or Fab (Fab1C06, Jiang et al.) blocking the luminal cavity by which SHH accesses the catalytic centre. In order to compare their binding mode, we determined the sequence and rebuilt the variable region of Fab1C06 using the deposited structure coordinates (PDB Id. 7MHY) and density (EMD-23836). This was necessary as part of the variable regions of Fab1C06 that interact with HHAT were unbuilt or built incorrectly as both the model and the attributed sequence did not match the experimental data (Fig. S11C and D), although this region is clearly one of the most ordered parts of the whole structure. For both MB177 and Fab1C06, one of the variable regions protrudes into the SHH-binding cavity but not deeply enough to impair binding of palm-CoA or affect organization of the catalytic centre (Fig. S11E). This suggests that their inhibition mechanism relies exclusively on competing with SHH binding which agrees with our competition binding experiment (Fig. 4E) and structural analysis (Fig. S8).

Structures in complex with either palm-CoA (Jiang et al., 2021) or nhPalm-CoA (this study) were obtained by adding excess during sample preparation. Both structures revealed the same conserved binding pocket with some small conformational changes in the binding mode near the catalytic centre that could be attributed to the chemical difference between the two molecules. However, we also observed a flip of the base in the CoA moiety and a rotation of the pantothenic moiety that are likely due to the quality of the data (Fig. S11F and G). Therefore, we further confirmed the conformation observed in our structures by a thorough structure-guided mutagenesis analysis (Fig. 3D and E). Jiang et al. also identify a density between TM10 and TM11 that they attributed to be a second palm-CoA referred as the “archway” palm-CoA. The density present in both cryo-EM maps (EMD-23836 & EMD-23837) is a non-continuous and has relatively weak signal compared to the density for the neighboring TM10 and TM11 (Fig. S11H). We also observe two discontinuous densities at a similar position and our data are in disagreement regarding the attribution of the density (Fig. S11I). The best fit for the density overlapping with the CoA moiety is an acyl chain that we attributed to be a palmitate based on the length of the density and hydrophobic environment. We also noticed that for both maps of Jiang et al, this part of the density is blurred due the intrinsic flexibility of the protein, complicating the interpretation of the map. For the density attributed to the “archway” Palm-CoA acyl tail, we also note that the size and position suggest that it belongs to the detergent (GDN) belt (Fig. S11I). Moreover, we have a clear, unambiguous density in our structure corresponding to the palmitoylation of Cys387 (Fig. S5, Fig. S11I), incompatible with binding of an “archway” Palm-CoA. To further probe the stability of the modeled “archway” palm-CoA, we performed atomistic MD simulations of the HHAT structure (5 x 100 ns) of Jiang *et al*. (PDB: 7mhy) bound to heme and with both palm-CoA molecules present (Fig. S11J). We observe a consistent displacement of the “archway” palm-CoA within the site by ∼9Å, which did not appear to be compatible with the coordinating residues proposed by Jiang *et al*. In addition, we observe dissociation of the “archway” palm-CoA tail away from HHAT in 4/5 replicates by 100 ns. We suggest the “archway” density observed by Jiang *et al*. could be accounted for by two lipid/detergent species in adjacent cytoplasmic and luminal leaflets, consistent with the break in continuous density observed towards the center of the modelled “archway” palm-CoA and with instabilities in palm-CoA binding at this site observed in our simulations.

Jiang *et al*. also determined the structure of HHAT bound to a palm-SHH peptide at low resolution (>3.2 Å); the connectivity of the palm-SHH peptide is broken, which limits interpretation of active site interactions (e.g. the SHH peptide carbonyl amide with active site HHAT His-379). However, the authors do observe a rearrangement of the active site residues, similar to those in our HHAT-IMP-1575 inhibitor complex (**Fig S11K**). IMP-1575 and analogues inhibit HHAT activity and paracrine Hh signalling in cells, and understanding the mechanism of action will assist further structure-guided development of potent and effective HHAT inhibitors. Importantly, the rearrangement of the active site in the HHAT-product (Jiang *et al*.) and inhibitor (Coupland *et al*.) complexes together with our BLI studies showing that the presence of the HHAT substrate palm-CoA enhances the SHH-HHAT interaction, suggest an ordered substrate binding mechanism. In the HHAT-product complex, Jiang et al show that HHAT-Glu59 as well as -Y382 and -Y384 possess a central role in the interaction with SHH. Our combined analyses using enzymatic assays (Fig. 3D and E) and binding studies with purified, monodisperse SHH and HHAT (Fig. 4F) confirms this finding, supporting the sequential mode of Palm-CoA and SHH binding to HHAT. Taking these structures together, it is plausible that CoA dissociates from the product complex before Palm-SHH, allowing rearrangement to release Palm-SHH and return apo HHAT, an order of events seen in other acyltransferases such as *N*-myristoyltransferase (Dian et al., 2020).

Our structural analysis also identified features additional to those reported by Jiang *et al*. For example, we identify an additional lipid density (modelled as a palmitate) that forms part of the HHAT palmitoyl-CoA binding site, thereby restricting the length of the acyl chain transferred to SHH (Fig. 2). We also observe another lipid binding pocket in juxtaposition to the luminal SHH-binding cavity (Fig. S9). The observed lipid density is suggestive of a sterol and could potentially act as a secondary binding site for SHH, by forming interactions with the C-terminally attached cholesteryl moiety of SHH.

Our additional studies combining MD simulations and mutagenesis analysis allow us to look beyond the protein structure towards HHAT mechanism and function in a cellular membrane environment. In particular, simulations of HHAT in a model ER membrane revealed substantial bilayer deformations that funnel towards the reaction centre (Fig. 2H **and** S7). This expands upon the single short simulation (90 ns) performed by Jiang *et al*. and emphasizes the importance of adequate sampling in simulations to observe changes in the configuration of the surrounding membrane environment. We suggest that HHAT-induced bilayer thinning may aid entry/exit of bound ligands. Also, in simulations we observe dynamic behaviour of the catalytic site Asp339 sidechain by which the two sidechain conformations are modulated by the absence or presence of bound palm-CoA. This agrees with the HHAT-product complex from Jiang *et al*., where Asp339 appears to stabilize the Cys24 amide nitrogen of the palm-SHH peptide.

## METHODS

### KEY RESOURCES TABLE

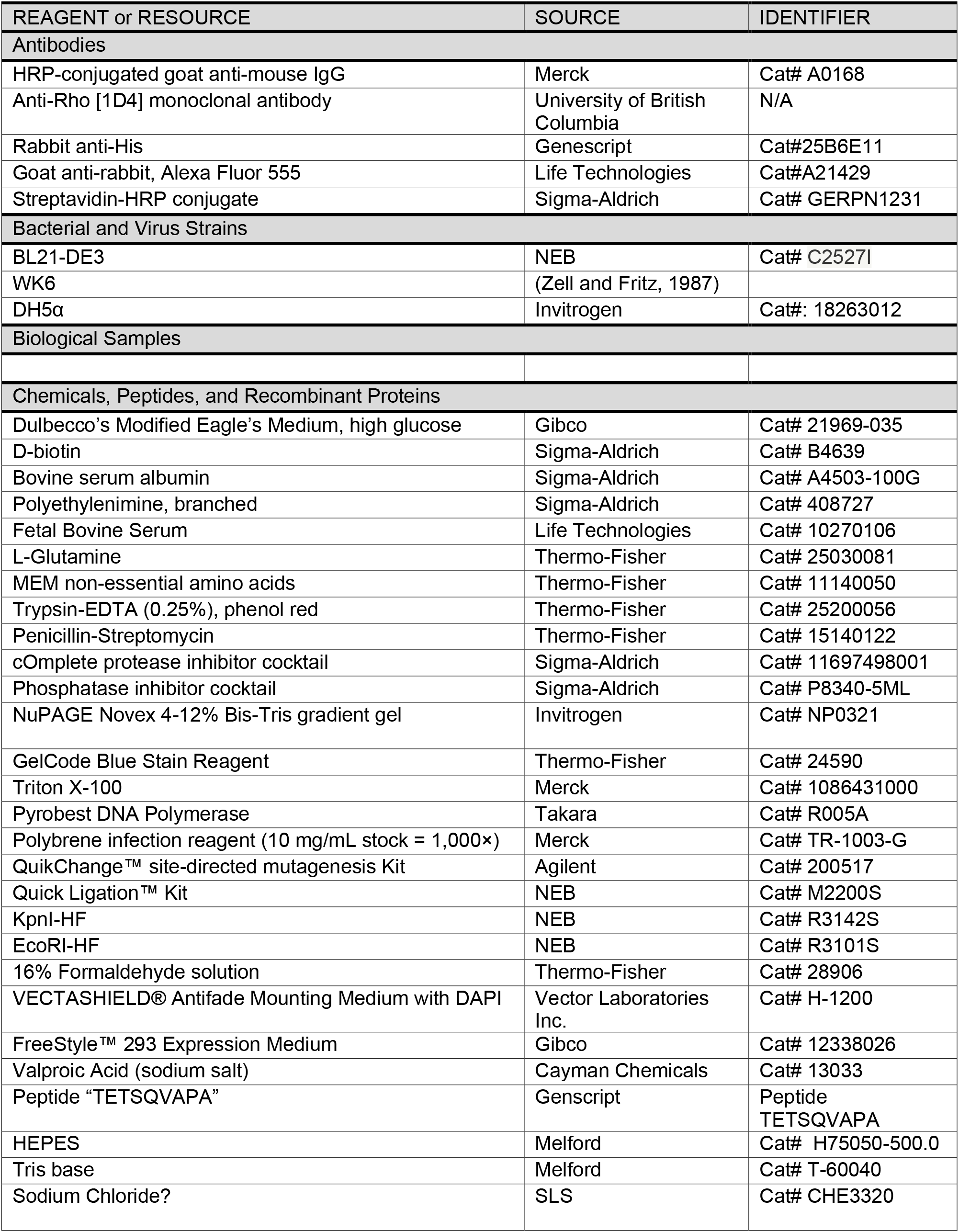
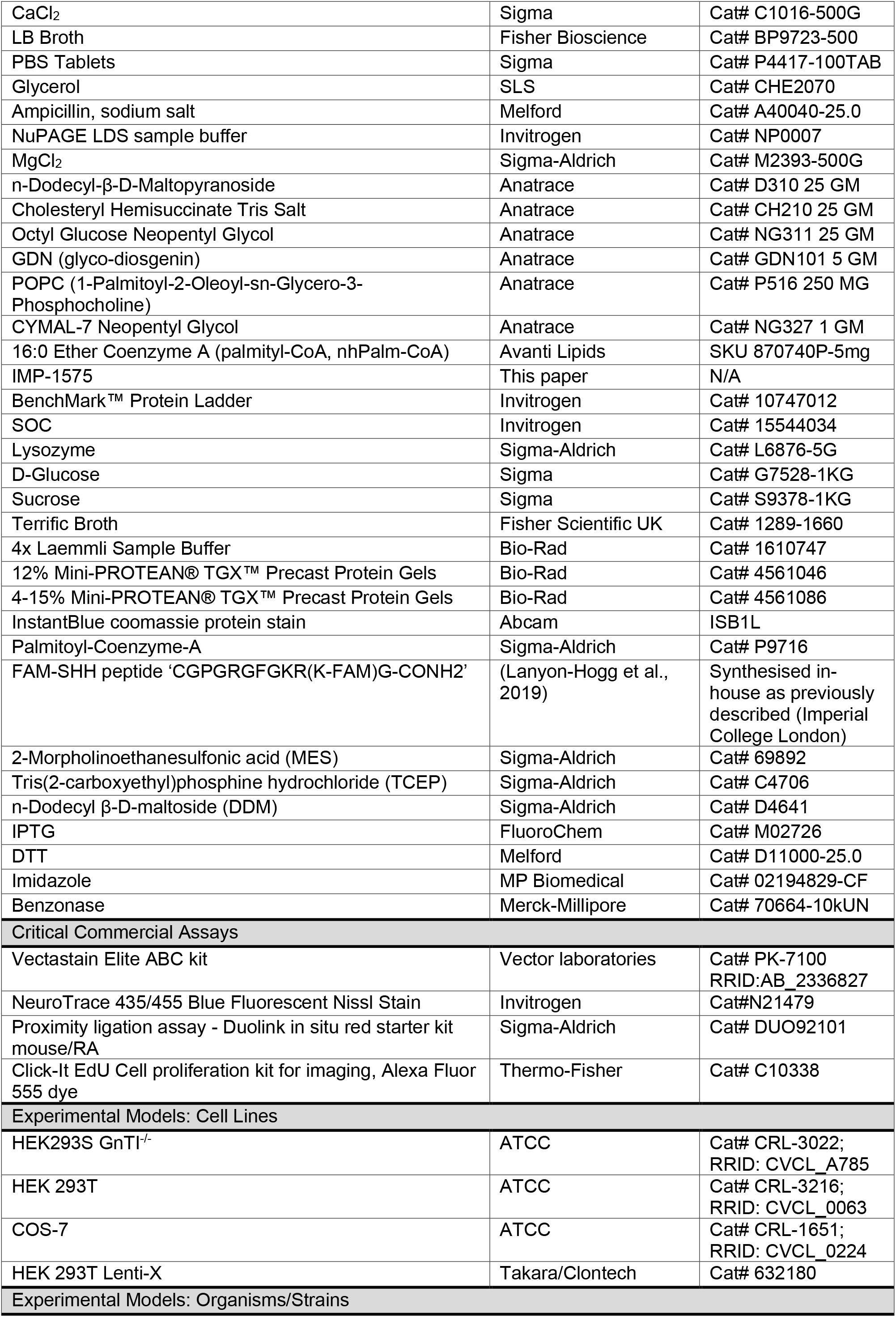
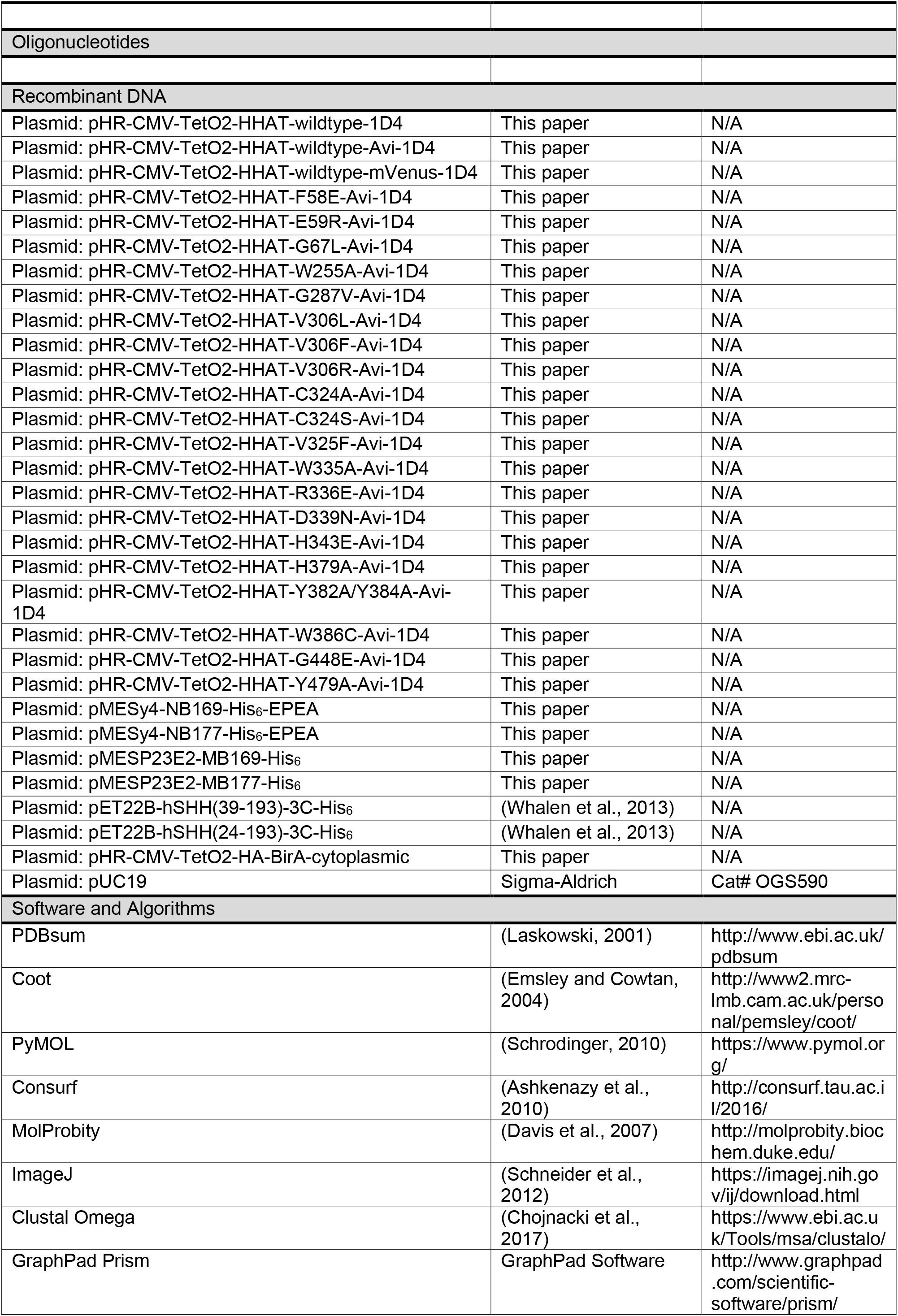
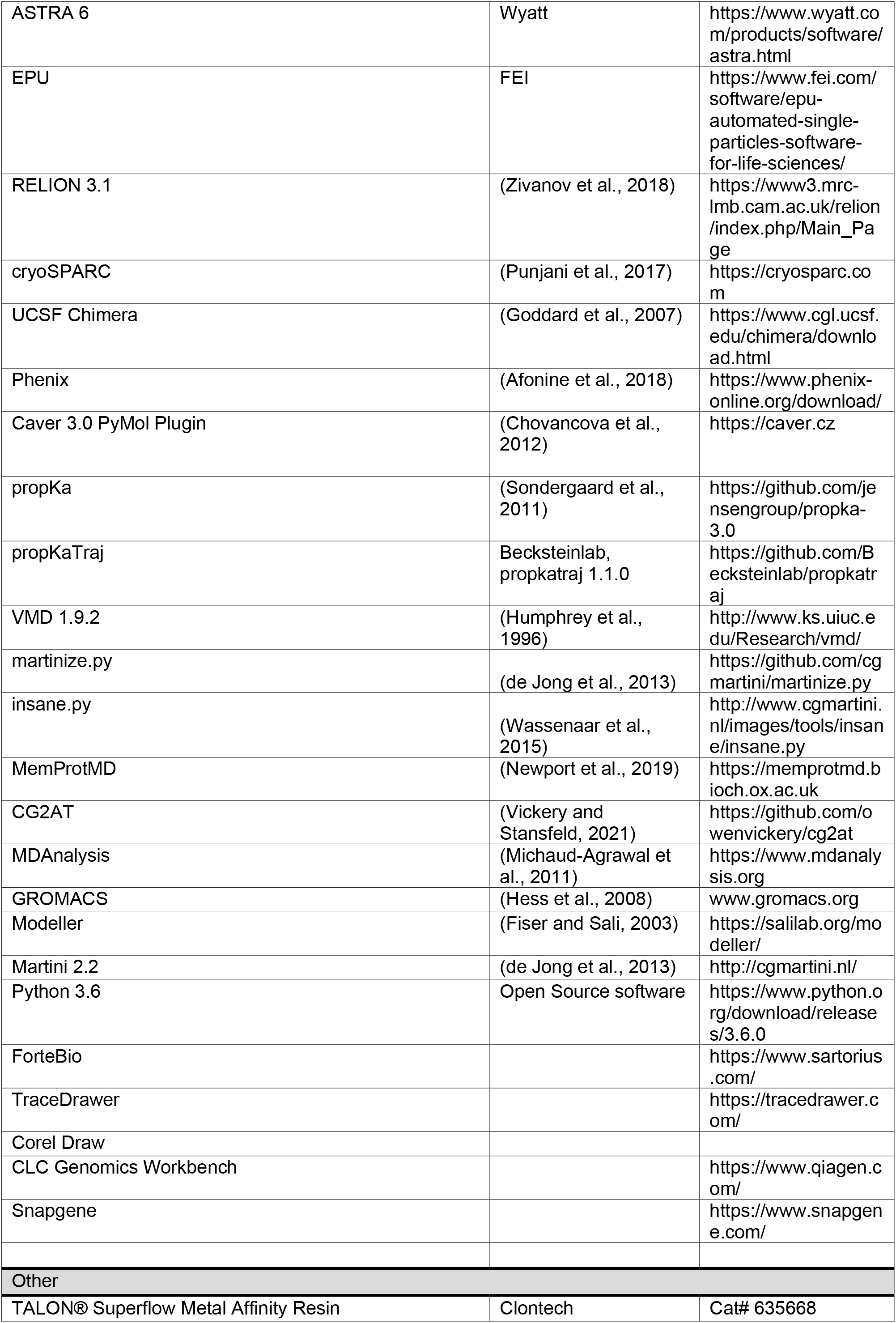
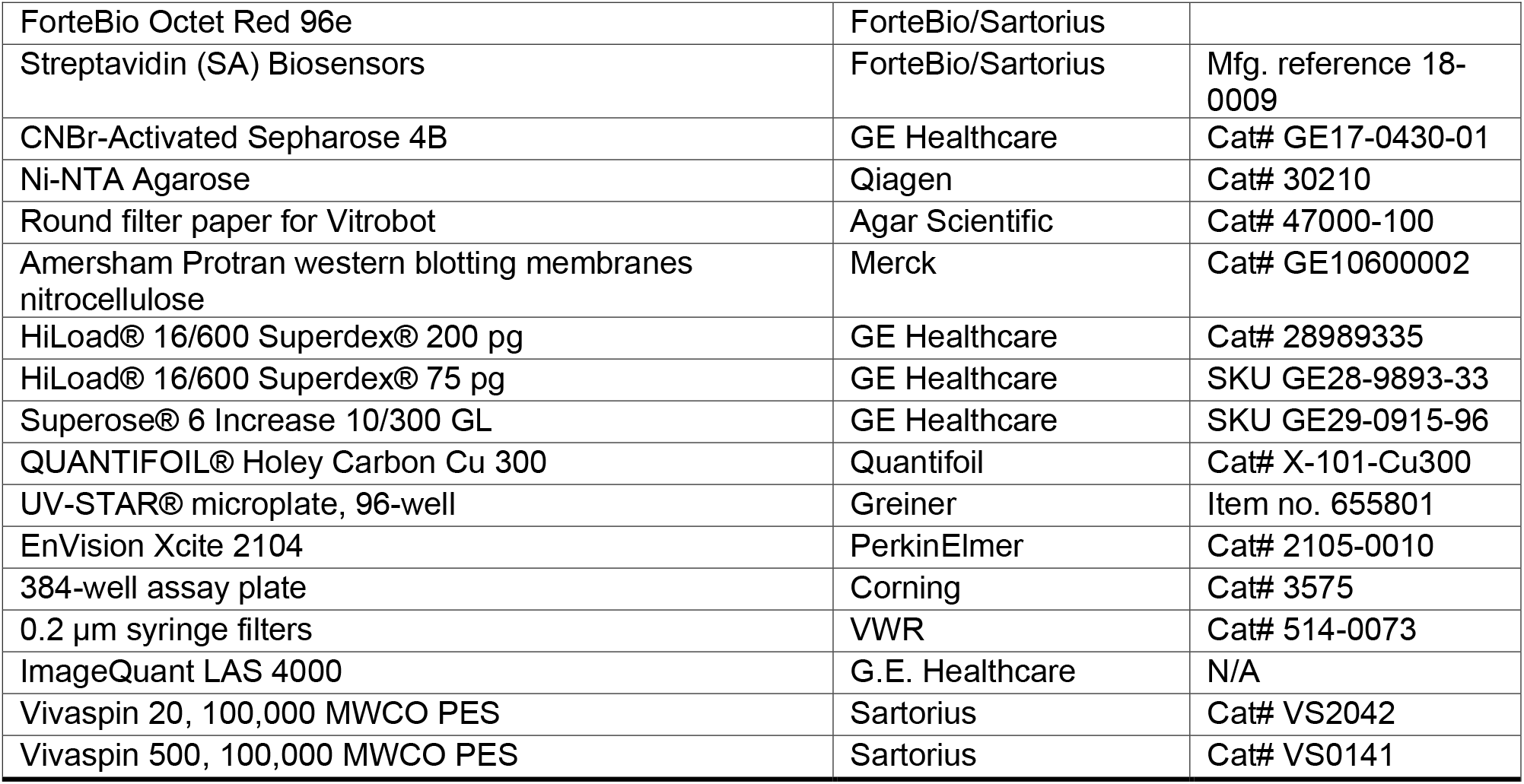

| REAGENT or RESOURCE | SOURCE | IDENTIFIER |
| --- | --- | --- |
| **Antibodies** |  |  |
| HRP-conjugated goat anti-mouse IgG | Merck | Cat# A0168 |
| Anti-Rho [1D4] monoclonal antibody | University of British Columbia | N/A |
| Rabbit anti-His | Genescript | Cat#25B6E11 |
| Goat anti-rabbit, Alexa Fluor 555 | Life Technologies | Cat#A21429 |
| Streptavidin-HRP conjugate | Sigma-Aldrich | Cat# GERPN1231 |
| **Bacterial and Virus Strains** |  |  |
| BL21-DE3 | NEB | Cat# C2527I |
| WK6 | (Zell and Fritz, 1987) |  |
| DH5α | Invitrogen | Cat#: 18263012 |
| **Biological Samples** |  |  |
| **Chemicals, Peptides, and Recombinant Proteins** |  |  |
| Dulbecco's Modified Eagle's Medium, high glucose | Gibco | Cat# 21969-035 |
| D-biotin | Sigma-Aldrich | Cat# B4639 |
| Bovine serum albumin | Sigma-Aldrich | Cat# A4503-100G |
| Polyethylenimine, branched | Sigma-Aldrich | Cat# 408727 |
| Fetal Bovine Serum | Life Technologies | Cat# 10270106 |
| L-Glutamine | Thermo-Fisher | Cat# 25030081 |
| MEM non-essential amino acids | Thermo-Fisher | Cat# 11140050 |
| Trypsin-EDTA (0.25%), phenol red | Thermo-Fisher | Cat# 25200056 |
| Penicillin-Streptomycin | Thermo-Fisher | Cat# 15140122 |
| cOmplete protease inhibitor cocktail | Sigma-Aldrich | Cat# 11697498001 |
| Phosphatase inhibitor cocktail | Sigma-Aldrich | Cat# P8340-5ML |
| NuPAGE Novex 4-12% Bis-Tris gradient gel | Invitrogen | Cat# NP0321 |
| GelCode Blue Stain Reagent | Thermo-Fisher | Cat# 24590 |
| Triton X-100 | Merck | Cat# 1086431000 |
| Pyrobest DNA Polymerase | Takara | Cat# R005A |
| Polybrene infection reagent (10 mg/mL stock = 1,000× | Merck | Cat# TR-1003-G |
| QuikChange™ site-directed mutagenesis Kit | Agilent | Cat# 200517 |
| Quick Ligation™ Kit | NEB | Cat# M2200S |
| KpnI-HF | NEB | Cat# R3142S |
| EcoRI-HF | NEB | Cat# R3101S |
| 16% Formaldehyde solution | Thermo-Fisher | Cat# 28906 |
| VECTASHIELD® Antifade Mounting Medium with DAPI | Vector Laboratories Inc. | Cat# H-1200 |
| FreeStyle™ 293 Expression Medium | Gibco | Cat# 12338026 |
| Valproic Acid (sodium salt) | Cayman Chemicals | Cat# 13033 |
| Peptide “TETSQVAPA” | Genscript | Peptide<br>TETSQVAPA |
| HEPES | Melford | Cat# H75050-500.0 |
| Tris base | Melford | Cat# T-60040 |
| Sodium Chloride? | SLS | Cat# CHE3320 |
| CaCl<sub>2</sub> | Sigma | Cat# C1016-500G |
| LB Broth | Fisher Bioscience | Cat# BP9723-500 |
| PBS Tablets | Sigma | Cat# P4417-100TAB |
| Glycerol | SLS | Cat# CHE2070 |
| Ampicillin, sodium salt | Melford | Cat# A40040-25.0 |
| NuPAGE LDS sample buffer | Invitrogen | Cat# NP0007 |
| MgCl<sub>2</sub> | Sigma-Aldrich | Cat# M2393-500G |
| n-Dodecyl-β-D-Maltopyranoside | Anatrace | Cat# D310 25 GM |
| Cholesteryl Hemisuccinate Tris Salt | Anatrace | Cat# CH210 25 GM |
| Octyl Glucose Neopentyl Glycol | Anatrace | Cat# NG311 25 GM |
| GDN (glyco-diosgenin) | Anatrace | Cat# GDN101 5 GM |
| POPC (1-Palmitoyl-2-Oleoyl-sn-Glycero-3-Phosphocholine) | Anatrace | Cat# P516 250 MG |
| CYMAL-7 Neopentyl Glycol | Anatrace | Cat# NG327 1 GM |
| 16:0 Ether Coenzyme A (palmitoyl-CoA, nhPalm-CoA) | Avanti Lipids | SKU 870740P-5mg |
| IMP-1575 | This paper | N/A |
| BenchMark™ Protein Ladder | Invitrogen | Cat# 10747012 |
| SOC | Invitrogen | Cat# 15544034 |
| Lysozyme | Sigma-Aldrich | Cat# L6876-5G |
| D-Glucose | Sigma | Cat# G7528-1KG |
| Sucrose | Sigma | Cat# S9378-1KG |
| Terrific Broth | Fisher Scientific UK | Cat# 1289-1660 |
| 4x Laemmli Sample Buffer | Bio-Rad | Cat# 1610747 |
| 12% Mini-PROTEAN® TGX™ Precast Protein Gels | Bio-Rad | Cat# 4561046 |
| 4-15% Mini-PROTEAN® TGX™ Precast Protein Gels | Bio-Rad | Cat# 4561086 |
| InstantBlue coomassie protein stain | Abcam | ISB1L |
| Palmitoyl-Coenzyme-A | Sigma-Aldrich | Cat# P9716 |
| FAM-SHH peptide 'CGPGRGFGKR(K-FAM)G-CONH2' | (Lanyon-Hogg et al., 2019) | Synthesised in-house as previously described (Imperial College London) |
| 2-Morpholinoethanesulfonic acid (MES) | Sigma-Aldrich | Cat# 69892 |
| Tris(2-carboxyethyl)phosphine hydrochloride (TCEP) | Sigma-Aldrich | Cat# C4706 |
| n-Dodecyl β-D-maltoside (DDM) | Sigma-Aldrich | Cat# D4641 |
| IPTG | FluoroChem | Cat# M02726 |
| DTT | Melford | Cat# D11000-25.0 |
| Imidazole | MP Biomedical | Cat# 02194829-CF |
| Benzonase | Merck-Millipore | Cat# 70664-10KUN |
| **Critical Commercial Assays** |  |  |
| Vectastain Elite ABC kit | Vector laboratories | Cat# PK-7100<br>RRID:AB_2336827 |
| NeuroTrace 435/455 Blue Fluorescent Nissl Stain | Invitrogen | Cat#N21479 |
| Proximity ligation assay - Duolink in situ red starter kit mouse/RA | Sigma-Aldrich | Cat# DUO92101 |
| Click-It EdU Cell proliferation kit for imaging, Alexa Fluor 555 dye | Thermo-Fisher | Cat# C10338 |
| **Experimental Models: Cell Lines** |  |  |
| HEK293S GnT1<sup>-/-</sup> | ATCC | Cat# CRL-3022;<br>RRID: CVCL_A785 |
| HEK 293T | ATCC | Cat# CRL-3216;<br>RRID: CVCL_0063 |
| COS-7 | ATCC | Cat# CRL-1651;<br>RRID: CVCL_0224 |
| HEK 293T Lenti-X | Takara/Clontech | Cat# 632180 |
| **Experimental Models: Organisms/Strains** |  |  |
| Oligonucleotides |  |  |
| Recombinant DNA |  |  |
| Plasmid: pHR-CMV-TetO2-HHAT-wildtype-1D4 | This paper | N/A |
| Plasmid: pHR-CMV-TetO2-HHAT-wildtype-Avi-1D4 | This paper | N/A |
| Plasmid: pHR-CMV-TetO2-HHAT-wildtype-mVenus-1D4 | This paper | N/A |
| Plasmid: pHR-CMV-TetO2-HHAT-F58E-Avi-1D4 | This paper | N/A |
| Plasmid: pHR-CMV-TetO2-HHAT-E59R-Avi-1D4 | This paper | N/A |
| Plasmid: pHR-CMV-TetO2-HHAT-G67L-Avi-1D4 | This paper | N/A |
| Plasmid: pHR-CMV-TetO2-HHAT-W255A-Avi-1D4 | This paper | N/A |
| Plasmid: pHR-CMV-TetO2-HHAT-G287V-Avi-1D4 | This paper | N/A |
| Plasmid: pHR-CMV-TetO2-HHAT-V306L-Avi-1D4 | This paper | N/A |
| Plasmid: pHR-CMV-TetO2-HHAT-V306F-Avi-1D4 | This paper | N/A |
| Plasmid: pHR-CMV-TetO2-HHAT-V306R-Avi-1D4 | This paper | N/A |
| Plasmid: pHR-CMV-TetO2-HHAT-C324A-Avi-1D4 | This paper | N/A |
| Plasmid: pHR-CMV-TetO2-HHAT-C324S-Avi-1D4 | This paper | N/A |
| Plasmid: pHR-CMV-TetO2-HHAT-V325F-Avi-1D4 | This paper | N/A |
| Plasmid: pHR-CMV-TetO2-HHAT-W335A-Avi-1D4 | This paper | N/A |
| Plasmid: pHR-CMV-TetO2-HHAT-R336E-Avi-1D4 | This paper | N/A |
| Plasmid: pHR-CMV-TetO2-HHAT-D339N-Avi-1D4 | This paper | N/A |
| Plasmid: pHR-CMV-TetO2-HHAT-H343E-Avi-1D4 | This paper | N/A |
| Plasmid: pHR-CMV-TetO2-HHAT-H379A-Avi-1D4 | This paper | N/A |
| Plasmid: pHR-CMV-TetO2-HHAT-Y382A/Y384A-Avi-1D4 | This paper | N/A |
| Plasmid: pHR-CMV-TetO2-HHAT-W386C-Avi-1D4 | This paper | N/A |
| Plasmid: pHR-CMV-TetO2-HHAT-G448E-Avi-1D4 | This paper | N/A |
| Plasmid: pHR-CMV-TetO2-HHAT-Y479A-Avi-1D4 | This paper | N/A |
| Plasmid: pMESy4-NB169-His<sub>6</sub>-EPEA | This paper | N/A |
| Plasmid: pMESy4-NB177-His<sub>6</sub>-EPEA | This paper | N/A |
| Plasmid: pMESP23E2-MB169-His<sub>6</sub> | This paper | N/A |
| Plasmid: pMESP23E2-MB177-His<sub>6</sub> | This paper | N/A |
| Plasmid: pET22B-hSHH(39-193)-3C-His<sub>6</sub> | (Whalen et al., 2013) | N/A |
| Plasmid: pET22B-hSHH(24-193)-3C-His<sub>6</sub> | (Whalen et al., 2013) | N/A |
| Plasmid: pHR-CMV-TetO2-HA-BirA-cytoplasmic | This paper | N/A |
| Plasmid: pUC19 | Sigma-Aldrich | Cat# OGS590 |
| Software and Algorithms |  |  |
| PDBsum | (Laskowski, 2001) | http://www.ebi.ac.uk/pdbsum |
| Coot | (Emsley and Cowtan, 2004) | http://www2.mrc-lmb.cam.ac.uk/personal/pemsley/coot/ |
| PyMOL | (Schrodinger, 2010) | https://www.pymol.org/ |
| Consurf | (Ashkenazy et al., 2010) | http://consurf.tau.ac.il/2016/ |
| MolProbity | (Davis et al., 2007) | http://molprobity.biochem.duke.edu/ |
| ImageJ | (Schneider et al., 2012) | https://imagej.nih.gov/ij/download.html |
| Clustal Omega | (Chojnacki et al., 2017) | https://www.ebi.ac.uk/Tools/msa/clustalo/ |
| GraphPad Prism | GraphPad Software | http://www.graphpad.com/scientific-software/prism/ |
| ASTRA 6 | Wyatt | https://www.wyatt.com/products/software/astra.html |
| EPU | FEI | https://www.fei.com/software/epu-automated-single-particles-software-for-life-sciences/ |
| RELION 3.1 | (Zivanov et al., 2018) | https://www3.mrc-lmb.cam.ac.uk/relion/index.php/Main_Page |
| cryoSPARC | (Punjani et al., 2017) | https://cryosparc.com |
| UCSF Chimera | (Goddard et al., 2007) | https://www.cgl.ucsf.edu/chimera/download.html |
| Phenix | (Afonine et al., 2018) | https://www.phenix-online.org/download/ |
| Caver 3.0 PyMol Plugin | (Chovancova et al., 2012) | https://caver.cz |
| propKa | (Sondergaard et al., 2011) | https://github.com/jensengroup/propka-3.0 |
| propKaTraj | Becksteinlab, propkatraj 1.1.0 | https://github.com/Becksteinlab/propkatraj |
| VMD 1.9.2 | (Humphrey et al., 1996) | http://www.ks.uiuc.edu/Research/vmd/ |
| martinize.py | (de Jong et al., 2013) | https://github.com/cgmartini/martinize.py |
| insane.py | (Wassenaar et al., 2015) | http://www.cgmartini.nl/images/tools/insane/insane.py |
| MemProtMD | (Newport et al., 2019) | https://memprotmd.bioch.ox.ac.uk |
| CG2AT | (Vickery and Stansfeld, 2021) | https://github.com/owenvickery/cg2at |
| MDAnalysis | (Michaud-Agrawal et al., 2011) | https://www.mdanalysis.org |
| GROMACS | (Hess et al., 2008) | www.gromacs.org |
| Modeller | (Fiser and Sali, 2003) | https://salilab.org/modeller/ |
| Martini 2.2 | (de Jong et al., 2013) | http://cgmartini.nl/ |
| Python 3.6 | Open Source software | https://www.python.org/download/releases/3.6.0 |
| ForteBio |  | https://www.sartorius.com/ |
| TraceDrawer |  | https://tracedrawer.com/ |
| Corel Draw |  |  |
| CLC Genomics Workbench |  | https://www.qiagen.com/ |
| Snapgene |  | https://www.snapgene.com/ |
| Other |  |  |
| TALON® Superflow Metal Affinity Resin | Clontech | Cat# 635668 |
| ForteBio Octet Red 96e | ForteBio/Sartorius |  |
| Streptavidin (SA) Biosensors | ForteBio/Sartorius | Mfg. reference 18-0009 |
| CNBr-Activated Sepharose 4B | GE Healthcare | Cat# GE17-0430-01 |
| Ni-NTA Agarose | Qiagen | Cat# 30210 |
| Round filter paper for Vitrobot | Agar Scientific | Cat# 47000-100 |
| Amersham Protran western blotting membranes nitrocellulose | Merck | Cat# GE10600002 |
| HiLoad® 16/600 Superdex® 200 pg | GE Healthcare | Cat# 28989335 |
| HiLoad® 16/600 Superdex® 75 pg | GE Healthcare | SKU GE28-9893-33 |
| Superose® 6 Increase 10/300 GL | GE Healthcare | SKU GE29-0915-96 |
| QUANTIFOIL® Holey Carbon Cu 300 | Quantifoil | Cat# X-101-Cu300 |
| UV-STAR® microplate, 96-well | Greiner | Item no. 655801 |
| EnVision Xcite 2104 | PerkinElmer | Cat# 2105-0010 |
| 384-well assay plate | Corning | Cat# 3575 |
| 0.2 µm syringe filters | VWR | Cat# 514-0073 |
| ImageQuant LAS 4000 | G.E. Healthcare | N/A |
| Vivaspin 20, 100,000 MWCO PES | Sartorius | Cat# VS2042 |
| Vivaspin 500, 100,000 MWCO PES | Sartorius | Cat# VS0141 |

### RESOURCE AVAILABILITY

#### Lead Contact

Further information and requests for reagents should be directed to the Lead Contact Christian Siebold.

#### Data and Code Availability

Cryo-EM density maps with the corresponding atomic coordinates will be deposited in the Electron Microscopy Data Bank with accession codes EMD-XXX (for the HHAT-MB177-nhPalm-CoA complex), EMD-YYY (for the HHAT-MB177-IMP1575 complex) and EMD-ZZZ (for the HHAT-MB169-nhPalm-CoA complex). Corresponding atomic coordinates for the HHAT-MB177-nhPalm-CoA, HHAT-MB177-IMP1575 and HHAT-MB169-nhPalm-CoA complexes will be deposited in the Protein Data Bank with accession codes XXX, YYY and ZZZ. Raw movies of the datasets for the HHAT-MB177-nhPalm-CoA, HHAT-MB177-IMP1575 and HHAT-MB169-nhPalm-CoA complexes will be deposited in the Electron Microscopy Public Image Archive (https://www.ebi.ac.uk/pdbe/emdb/empiar/) with accession codes EMPIAR-XXX, EMPIAR-YYY and EMPIAR-ZZZ, respectively.

### EXPERIMENTAL MODEL AND SUBJECT DETAILS

#### Cell lines

All cell lines used in this study are listed in the Key Resources Table and were cultured under standard growth conditions (37 °C, 5% CO_2_). HEK293T were used for transient mammalian expression using pHLsec vector (Aricescu et al., 2006). HEK293T Lenti-X cells were utilised to generate lentiviruses with the pHR-CMV-TetO_2_ vector (Elegheert et al., 2018a), which were subsequently used to infect HEK293T cells and produce stable cell lines. Both cell lines were maintained in Dulbecco’s Modified Eagle’s Medium (DMEM high glucose, Gibco) supplemented with L-glutamine, MEM non-essential amino-acids (NEAA; both Gibco) and 10% fetal bovine serum (FBS, Life Technologies) with no addition of antibiotics. Cells were grown and maintained in standard T75 (75 cm^2^ - LentiX) or T175 (175 cm^2^ - HEK293T) flasks. Cos-7 cells were used for immunofluorescence staining experiments (transfected with pHR-CMV-TetO_2_ vectors). Cos-7 cells were grown in complete medium consisting of DMEM supplemented with 10% (v/v) FBS, L-Glutamine and NEAA. For immunofluorescence staining experiments, Cos-7 cells were incubated with nanobodies in complete medium. We used *E. coli* DH5α competent cells for cloning and *E. coli* BL21-DE3 for expression of ShhN. *E. coli* WK6 cells were used for nanobody and megabody expression.

### METHOD DETAILS

#### HHAT expression and purification

For protein expression, DNA encoding wild-type HHAT from *Homo sapiens* (VAR_024743; Uniprot ID: Q5VTY9) was cloned into pHR-CMV-TetO_2_ vector (Elegheert et al., 2018b) in frame with a C-terminal Rho1D4 antibody epitope tag (Molday and MacKenzie, 1983) (TETSQVAPA) alone (pHR-CMV-TetO2-HHAT-1D4), preceded by monoVenus (pHR-CMV-TetO2-HHAT-mVenus-1D4) or biotinylation tag (GLNDIFEAQKIEWHE) (pHR-CMV-TetO2-HHAT-Avi-1D4). For production of HHAT mutants, the HHAT gene was cloned into the pUC19 cloning plasmid and point mutations were introduced by QuikChange™ site-directed mutagenesis (Agilent) and then transferred into pHR-HHAT-Avi-1D4.

For expression, stable cell lines were created as described (Elegheert et al., 2018b). HEK293S Lenti-X producer cells (Takara/Clontech) were transfected in DMEM with 2 % (v/v) FBS with 12 μg of pHR-CMV-TetO2 construct vector alongside 11 μg envelope plasmid (pMD2.G; Addgene #12259) and 11 μg packaging plasmid (psPAX2; Addgene #12260) using PEI at 2.5:1 (w/w) ratio to total DNA. After 2 days cell culture media containing viral particles was filtered, supplemented with 1 μg/mL polybrene and added to HEK293S GnTI-TetR cells. For biotinylated samples, media containing viral particles for HHAT constructs was mixed in a 50:1 (v/v) ratio with media from Lenti-X cells transfected as above with biotin ligase (BirA) in a pHR-CMV-TetO_2_ vector and then added to HEK293S cells. These cells were incubated 37 °C, 8 % CO_2_ for at least 2 passages before expansion or freezing and transfer to cryo-storage. Stably-transfected HEK293S cell cultures were grown in 2-10 L suspension at 37 °C, 130 rpm, 8 % CO_2_ in Freestyle 293 media, supplemented with 1 % (v/v) FBS, 1 % L-glutamine and NEAA (Gibco) for at least 2 passages. Cells at 2-3 x 10^6^ cells/mL were induced with 50 ng/mL doxycycline (Merck) and 5 mM valproic acid (Cayman Chemical). For biotinylated proteins, 0.8 mM biotin was added the day preceding induction. Media was harvested after 72 h, centrifuged (Beckman Avanti, 1500 x g, 10 min, 4 °C) and pellets were flash-frozen and stored at −80 °C.

For purification of HHAT for cryo-EM, all steps were performed at 4 °C. Cells were lysed and membranes solubilised in 20 mM Tris pH 8.0, 300 mM NaCl, 5 % (v/v) glycerol, 1 mM phenylmethylsulfonyl fluoride (PMSF), with 1 % (v/v) membrane protease inhibitors (Sigma-Aldrich) and 1.5 % (w/v) Octyl Glucose Neopentyl Glycol (OGNG) 40:1 (w/w) cholesteryl hemisuccinate (CHS) (Anatrace) by 90 min stirring. Cell debris was removed by centrifugation (Beckman Avanti, 45 min, 40,000 x g, 4 °C) and supernatants were diluted 3-fold in 20 mM Tris pH 8.0, 300 mM NaCl, 5 % (v/v) glycerol, before incubation 2 h with 500 μL CNBr-activated Sepharose beads (GE Healthcare) coupled to monoclonal Rho-1D4 antibody (University of British Columbia) per 1 g pellet. Beads were collected on a flow column and washed with 20 column volumes (CV) 20 mM Tris pH 8.0, 300 mM NaCl (wash buffer, WB) with 0.522 % (w/v) OGNG 40:1 CHS, then 20 CV WB with 0.174 % (w/v) OGNG 40:1 CHS, 20 CV WB with 0.06 % (w/v) glyco-diosgenin (GDN) (Anatrace) and 20 CV WB with 0.02 % (w/v) GDN. HHAT was eluted by overnight incubation with 500 μM 1D4 (TETSQVAPA) peptide (Genscript).

Purified megabody was added at 1:1.5 molar ratio and 0.05 % (w/v) CYMAL-7-NG (Anatrace) and 0.01 mg/mL 1-palmitoyl-2-oleoyl-sn-glycero-3-phosphocholine (POPC) were added before concentration in Vivaspin® 100,000 kDa molecular weight cut-off (MWCO) centrifugal concentrators (Sartorius) to a volume of ∼500 μL. The sample was passed over a Superose 6 Increase 10300 GL column (GE Healthcare), equilibrated in 10 mM Tris pH 8.0, 150 mM NaCl, 0.02 % (w/v) GDN. Protein-containing fractions were pooled and concentrated to 4 mg/mL.

Biotinylated HHAT and HHAT mutants were purified in a similar manner to above. Membranes were solubilised in 20 mM HEPES pH 7.5, 300 mM NaCl, 5 % (v/v) glycerol, 1 mM PMSF, with 1 % (v/v) membrane protease inhibitors and 1.0 % (w/v) n-dodecyl-β-D-maltopyranoside (DDM) 40:1 (w/w) CHS (Anatrace). After centrifugation, supernatants were diluted 2-fold in 20 mM HEPES pH 7.5, 300 mM NaCl, 5 % (v/v) glycerol, before incubation 2 h with 100 μL Rho-1D4 antibody-coupled Sepharose beads per 1 g pellet. Beads were washed with 25 column volumes (CV) 20 mM HEPES pH 7.5, 300 mM NaCl (wash buffer, WB) with 0.09 % (w/v) DDM 40:1 CHS, then 75 CV WB with 0.03 % (w/v) DDM 40:1 CHS. HHAT was eluted by overnight incubation with 500 μM 1D4 (TETSQVAPA) peptide (Genscript). Protein was flash-frozen and stored at −80 °C before direct use in biolayer interferometry or enzyme activity assays.

For HHAT samples for SEC-MALLS, HHAT was purified in a similar manner to the biotinylated samples from cells expressing the pHR-HHAT-mVenus-1D4 construct. Membranes were solubilised in 20 mM HEPES pH 7.5, 300 mM NaCl, 5 % (v/v) glycerol, 1 mM ethylenediaminetetraacetic acid (EDTA), 10 mM L-arginine with 0.5 % (v/v) membrane protease inhibitors (Sigma-Aldrich) and 1.5 % (w/v) OGNG 40:1 (w/w) CHS. Rho-1D4 antibody-coupled Sepharose beads were added after centrifugation, as above and washed with 50 column volumes (CV) 20 mM HEPES pH 7.5, 300 mM NaCl (wash buffer, WB) with 0.580 % (w/v) OGNG 40:1 CHS, then 50 CV WB with 0.174 % (w/v) OGNG 40:1 CHS. HHAT was eluted by overnight incubation with 3C protease. Elution was concentrated in Vivaspin® 100,000 kDa molecular weight cut-off (MWCO) centrifugal concentrators (Sartorius) to a volume of ∼800 μL. The sample was passed over a Superose 6 Increase 10300 GL column (GE Healthcare), equilibrated in 10 mM HEPES pH 7.5, 150 mM NaCl, 0.174 % (w/v) OGNG 40:1 CHS. Protein-containing fractions were pooled and concentrated to 1 mg/mL.

#### Generation of nanobodies and megabodies

Nanobodies were constructed as described in (Pardon et al., 2014). cDNA derived from Llama (*Lama glama*) serum was cloned into the pMESy4 (GenBank KF415192) phage display and bacterial expression vector. Megabody-expressing vectors were constructed as described in (Uchanski et al., 2021) via insertion of nanobody-encoding DNA into pMESP23E2 (GenBank: MT338521.1), which contains a circular permutant of *Escherichia coli* K12 glucosidase YgjK (Uniprot P42592; Gene ID: 947596) inserted into the first β-turn of Nb177.

Nanobodies and megabodies with C-terminal His_6_ tags were expressed in WK6 *E. coli* cells grown to an optical density of 0.7-1.2 in terrific broth supplemented with ampicillin at 37 °C, 180 rpm. Expression was induced with 1 mM isopropyl β-d-1-thiogalactopyranoside (IPTG) and incubation temperature was lowered to 20 °C overnight. Periplasmic extraction was performed in PE buffer (50 mM Tris pH 8.0, 150 mM NaCl, 20 % (w/v) sucrose, 0.4 mg/mL lysozyme, DNase, 1 mM PMSF, protease inhibitor cocktail) and incubated at 4 °C for 30 min with shaking. The soluble fraction was removed by centrifugation. Proteins were purified by nickel-affinity chromatography and samples were passed over a HiLoad® 16/600 Superdex® 75 or 200 pg column (GE Healthcare), equilibrated in 20 mM Tris pH 8.0, 300 mM NaCl. Protein-containing fractions were concentrated to 50-400 μM, before addition to HHAT to form complexes, except for the HHAT-IMP-1575-MB177 complex, where purified MB177 was flash-frozen and stored at −80 °C.

#### Sonic Hedgehog (SHH) expression and purification

Wild-type sonic hedgehog (SHH) from *H. sapiens* (Uniprot ID: Q15465, residues 24-193) was cloned into the pET22B bacterial expression vector, with a C-terminal His_6_ tag. SHH was expressed from *E. coli* Rosetta (DE3) pLysS cells using an adapted protocol described previously (Whalen et al., 2013). Cultures were grown in LB at 37 °C, 180 rpm to an optical density of 0.6-0.8 before induction with 1 mM IPTG and decrease of temperature to 20 °C overnight. Harvested cell pellets were resuspended in (1 x PBS, + 21 g NaCl, + 70 μL beta-mercaptoethanol, 1 mM PMSF, 1 protease inhibitor tablet). Cells were lysed using a cell disruptor and cell debris was removed via centrifugation. The protein was purified from the supernatant using TALON® beads. The sample was then further purified using size exclusion chromatography (SEC) (HiLoad® 16/600 Superdex® 75 pgcolumn, GE Healthcare) equilibrated in 5 mM Na_2_HPO_4_ pH 5.5, 150 mM NaCl, 0.5 mM dithiothreitol (DTT)). Protein-containing fractions were concentrated and snap-frozen in liquid nitrogen for storage at −80 °C.

#### Cryo-EM sample preparation, image collection and processing

Purified HHAT protein was mixed with the corresponding purified megabody at 1:1.5 molar ratio and purified by SEC using a Superose 6 Increase 10/300 GL column in 20 mM Tris-HCl, pH 8.0, 150 mM NaCl, at 4 °C. The purified complex was concentrated to ∼ 4 mg/mL. For ligand-bound complexes, a final concentration of 5 mM nhPalm-CoA or 1 mM IMP-1575 inhibitor was added to the solution before grid plunging. A sample volume of 3.5 μl was placed on glow-discharged 300 copper mesh Quantifoil Holey Carbon grids R 1.2/1.3, before blotting for 3.0 sec and flash-freezing in liquid ethane. All grids were prepared using a Vitrobot mark IV (FEI) at 4 °Ç 95-100 % humidity.

Cryo-EM data for the HHAT-MB177-nhPalm-CoA, HHAT-MB169-nhPalm-CoA and HHAT-MB177-IMP-1575 sample were collected on a 300 kV Titan Krios microscope (Thermo Fisher Scientific) at the Electron Bio-Imaging Centre (eBIC). Automated data collection was setup in EPU using a K3 (Gatan) direct electron detector operating in super-resolution mode with a physical pxel size of 0.829 Å/px (0.4145Å/px super-resolution pixel) or 0.831 Å/px (0.4155Å/px super-resolution pixel and a GIF Quantum energy filter (Gatan) with 20 eV slit. Sample was collected with a total dose of ∼ 53-61 e-/Å2 across 50 frames. Sample-specific data collection parameters are summarized in Table S1.

Data processing pipelines are shown in Fig. S2. Briefly, data were processed using cryoSPARC v.3.1.1 (Punjani et al., 2017) standard workflow. Raw movies were aligned with patch motion correction and binned 2 times at the motion correction step, giving a final pixel size of 0.829 Å/px or 0.831 Å/px. The contrast transfer function (CTF) was initially estimated using Patch-CTF. Poor-quality images were discarded after manual inspection. Particles were blob picked and the 2D classes were inspected and classes of interest were selected to generate templates for complete particle picking.

For the HHAT-MB177-nhPalm-CoA complex, a total number of 606,945 particles were picked and extracted in a 400 px box Fourier cropped to 200 px (1.662 Å/px). After template picking, 2D classification was skipped and 5 ab initio models were directly generated and further refined using heterogenous refinement, leading to a single good-looking class. Particles within this class having a 3D ab initio class posterior probability lower than 90% were discarded (39,331 particles) and the other particles were recentred and re-extracted in a 400 px box Fourier cropped to 300 px (1.1053 Å/px) before running a non-uniform refinement. The consensus map was refined to 3.0 Å. CTF refinement and high order aberration correction improved map resolution to 2.89 Å. Particles were exported to Relion 3.1.1 (Zivanov et al., 2018) using csparc2star.py and Bayesian polished (Asarnow et al., 2019). Focus refinement in cryoSPARC after particles subtraction was performed on the nanobody-HHAT part improving the resolution to 2.7 Å. A round of 3D classification without alignment in Relion with 10 classes was performed, leading to a major class with 238k particles. This class was further refined using focus refinement in cryoSPARC, and despite the same reported resolution, the quality of the map was significantly improved. In parallel, the particles have been reverted to the original and a focus refinement after particles subtraction has been performed on the megabody core, leading to a resolution of 2.6 Å. Both focus refined maps have been combined using Phenix combine focused map followed by phenix autosharpen map tool (Adams et al., 2010). The combined map has been used for model building after assessing that the quality of the combined map was identical to the focused maps.

For the HHAT-MB169-nhPalm-CoA complex, a total number of 2,300,000 particles were picked and extracted in a 400 px box Fourier cropped to 200 px (1.662 Å/px). Around 1,000,000 particles were selected after 2D classification and 5 ab initio models were then generated and further refined using heterogenous refinement, leading to a single good-looking class. Particles within this class having a 3D ab initio class posterior probability lower than 90% were discarded and the other particles were recentred and re-extracted in a 400 px box Fourier cropped to 300 px (1.1053 Å/px) before running a non-uniform refinement. The consensus map was refined to 3.4 Å. Particles were exported to Relion 3.1.1 (Zivanov et al., 2018) using csparc2star.py (Asarnow et al., 2019) and Bayesian polished before an extra step of 3D classification. The best-looking class containing 192K particles was further refined to 3.1 Å. Focus refinement on the nanobody-HHAT part greatly improved the quality of the transmembrane regions of HHAT with a final resolution of 3.74 Å.

Finally, for the HHAT-MB177-IMP-1575 complex, a total number of around 3,000,000 particles were picked and extracted in a 400 px box Fourier cropped to 200 px (1.62 Å/px). Around 864k particles were selected after 2D classification and 5 ab initio models were then generated and further refined using heterogeneous refinement, leading to two good-looking class. Particles within these classes having a 3D ab initio class posterior probability lower than 90% were discarded and the other particles were recentred and re-extracted in a 400 px box Fourier cropped to 300 px (1.1053 Å/px) before running a non-uniform refinement. The consensus map containing around 367k particles was refined to 4.0 Å. After an extra step of 3D classification, the best-looking class containing 318K particles was further refined to 3.86 Å. Focus refinement on the nanobody-HHAT part greatly improved the quality of the transmembrane regions of HHAT with a final resolution of 3.59 Å.

#### Model building, refinement and validation

The structures were modelled by first fitting homology model of the HHAT and pdb 6XUX for the megabody component into the combined map of HHAT-MB177-nhPalm-CoA using UCSF Chimera (Pettersen et al., 2004). One cycle of rigid body real space refinement followed by manual adjustment in Coot (Emsley et al., 2010) was performed to correctly position the Cα chain into the density. Finally, cycles of Phenix. (Adams et al., 2010) real space refinement and manual building in Coot (Emsley et al., 2010) were used to improve model geometry. Map-to-model comparison in Phenix. mtriage validated that no over-fitting was present in the structures. Model geometry was validated for all models using MolProbity (Chen et al., 2010). This model was then used to refine the HHAT-MB169-nhPalm-CoA and HHAT-MB177-IMP-1575 complexes in a similar way. All map and model statistics are detailed in Table S1.

#### Coarse-grained molecular dynamics simulations

Ligands and the bound megabody were removed from the HHAT-MB177-nhPalm-CoA-heme structure and loops between D45-L51, W189-T197 and P409-Q422 modelled using Modeller 9.20 (salilab.org/modeller). The model with the lowest DOPE score was selected and converted to CG resolution using *matinize.py*. The MARTINI 2.2 forcefield (de Jong et al., 2013) was used to describe all components and the ElNeDyn elastic network applied to HHAT (force constant: 1000 kJ mol^-1^ nm^-2^, cut-off: 0.9 nm) (Periole et al., 2009). HHAT was positioned in a bilayer composed of POPC (35%), DOPC (35%), POPE (8%), DOPE (7%), cholesterol (10%) and palmitate (5%) in the luminal leaflet and POPC (15%), DOPC (15%), POPE (19%), DOPE (18%), POPS (8%), PIP_2_ (10%), cholesterol (10%) and palmitate (5%) in the cytoplasmic leaflet using *insane.*py (Wassenaar et al., 2015). This bilayer composition was selected to recapitulate the main features of ER membranes e.g., low sphingolipid content and a cholesterol:phospholipid ratio of approximately 0.15 (Jacquemyn et al., 2017). Palmitate was modelled with a charged headgroup (named PCN in the MARTINI topology) and cholesterol was modelled with inclusion of virtual sites. The membrane position around DltB (PDB: 6BUG) on the MemProtMD server (memprotmd.bioch.ox.ac.uk) was used to guide HHAT insertion into the bilayer. The system was solvated using *insane.py* (Wassenaar et al., 2015) and approximately 0.15 M NaCl added to yield a total box size of 14 x 14 x 9 nm^3^. Prior to the production run, each replicate was independently energy minimised using a steepest decent method and equilibrated in two steps (step 1: 25 ns with all protein beads restrained, step 2: 100 ns with backbone beads restrained).

CG MD simulations (10x 15 μs) were performed using GROMACS 2019.4 (www.gromacs.org). The timestep was 20 fs and periodic boundary conditions were applied. Coulombic interactions were described using the reaction-field method and a 1.1 nm cut-off. Van der Waals interactions were described with the potential-shift Verlet method and a 1.1 nm cut-off. The V-rescale thermostat (Bussi et al., 2007) was used to maintain temperature as 310 K with a 1.0 ps coupling time constant. Pressure was maintained at 1 bar using the Parrinello-Rahman barostat, a 12.0 ps coupling time and a compressibility of 3x 10^-4^ bar^-1^.

#### Atomistic molecular dynamics simulations

A snapshot from the CG simulations was selected for backmapping to atomistic resolution using a revised CG2AT protocol (Vickery and Stansfeld, 2021) (github.com/owenvickery/cg2at). Prior to backmapping, palmitate was removed from the bilayer and a short equilibration performed to close gaps in the bilayer since topology files for backmapping palmitate were not available. The protein conformation was backmapped to the structural coordinates such that only the lipid arrangement reflected those of the CG snapshot. TIP3P water was readded to the system along with approximately 0.15 M NaCl.

The protein coordinates from the HHAT-nhPalm-CoA-heme structure were used to build the following systems: HHAT apo, HHAT with Palm-CoA bound and HHAT with Palm-CoA bound and heme covalently bound to Cys324. The protein coordinates from the HHAT-IMP-1575 structure were used to build the following system: HHAT in an inhibited conformation (but without inhibitor bound) with heme covalently bound to Cys324. The protonation state of residue sidechains in the input structures were assessed using propKa (github.com/jensengroup/propka-3.0) which did not identify any unusual pKas, therefore D339 was modelled as deprotonated and H379 was neutral. The CHARMM-36 forcefield (Huang and MacKerell, 2013) was used to describe all components. Palm-CoA parameters were derived from Stix *et al* (Stix et al., 2020). Standard CHARMM-36 parameters were used for the Mg ion bound to Palm-CoA (which remained bound without the need for additional restraints) and for the heme/Fe complex. To describe heme-Cys324 bonding Cys324 was modelled as a thiolate (named CYM in the CHARMM-36 forcefield) and S-Fe bond parameters described identically to existing parameters for heme coordinated by methionine. Control simulations of heme without bonding to Cys324 were also performed, yielding similar results (data not shown). Each replicate was energy minimised using a steepest decent method then subject to 2x 5 ns NVT and NPT equilibration steps with position restraint applied to the protein backbone and the heavy atoms of bound ligands.

Each system was simulated for 5x 200 ns with a 2 fs timestep using GROMACS 2019.4 (www.gromacs.org). The Particle-Mesh Ewald (PME) method was used for long-range electrostatic interactions with a 1.2 nm cut-off. Van der Waals interactions were smoothly switched between 1.0 and 1.2 nm using the force-shift modifier. Temperature was maintained at 310 K using the Nosé-Hoover thermostat with a 0.5 ps coupling time. Pressure was maintained at 1 bar using the Parrinello-Rahman barostat, a 2.0 ps coupling time and a compressibility of 4.5 x 10^-5^ bar^-1^. A dispersion correction was not applied. The LINCS algorithm was used to constrain bonds to their equilibrium values.

In simulations initiated from the HHAT apo conformation a DOPC lipid bound within the luminal gate was removed after 200 ns in each of the 5 replicates. After lipid removal, each replicate was simulated for an additional 200 ns.

#### Steered molecular dynamics simulations

In addition to removal of the DOPC lipid (at t = 200 ns) from the luminal gate (described above) atomistic simulations were also performed where the lipid was pulled laterally into the membrane over a period of 10 ns before 40 ns of additional unbiased simulation time for each of the five replicates. Steered MD simulations were assisted by use of PLUMED (consortium, 2019). The collective variable used during steered MD was defined as the distance between the centre of mass of the DOPC lipid and the Cα atom of Arg176. A moving restraint was applied over 10 ns to gradually increase the force constant to k=1000 kJ mol^-1^ nm^-2^. This restraint was switched back to k=0 kJ mol^-1^ nm^-2^ over the following 0.5 ns before a further 39.5 ns of unbiased simulation. All other simulations conditions are identical to those described above.

#### Simulation analysis

MDAnalysis (Michaud-Agrawal et al., 2011) (www.mdanalysis.org) was used for solvent and lipid density calculations. PropKaTraj (github.com/Becksteinlab/propkatraj) was used to calculate pKa values over the trajectory. PyMol (https://pymol.org) and VMD (Humphrey et al., 1996) were used for visualisation. Tunnels were calculated using the Caver3.0 plugin (https://caver.cz) implemented in PyMol.

#### Detection of HHAT binding to nanobodies NB169 and NB177

Cos-7 cells were infected with pHR-HHAT-mVenus-1D4, expressing full-length human HHAT C-terminally fused to mono-Venus. Once the stable cell line was established, cells were plated onto glass coverslips and incubated for 24 hours at 37 °C and 5 % CO_2_. All subsequent procedures were carried out at room temperature. Initially, cells were washed with PBS and fixed in 4% paraformaldehyde for 10 min. Fixed cells were then permeabilised with 0.1 % Triton X-100/PBS for 10 min and blocked with 10 % FBS/PBS for 1h. Where indicated, cells were incubated with 2.5 μM nanobody in PBS for 1h. This was followed by several PBS wash steps and incubation with anti-His antibody (#25B6E11, Genscript) at a dilution of 1:500 for 1h in 0.1 % FBS/PBS. Unbound primary antibody was removed with PBS washes and bound His antibody was recognised with an anti-rabbit AF555 antibody incubated for 1h in 0.1% FBS/PBS. Final wash steps were followed by mounting the cover slips on microscopy slides using Vectashield with Dapi. Slides were viewed on a Leica DMi8 microscope with a 100x oil immersion objective (HC PL APO, NA 1.47) and a Leica EL6000 fluorescence illuminator. Images were taken with a Hamamatsu Orca Flash 4.0 V2 Camera and processed in Fiji ImageJ (Schindelin et al., 2012).

#### Size exclusion chromatography coupled with multi angle light scattering (SEC-MALS)

For SEC-MALS, OGNG-solubilised HHAT at 1 mg ml^−1^ was loaded onto a Superose 6 10/300 GL column (GE Healthcare), equilibrated in 10 mM HEPES pH 7.5, 150 mM NaCl, 0.174 % OGNG 40:1 CHS, on a Shimadzu system with an inline Dawn HELEOS-II 8-angle light scattering detector (Wyatt). Scattering data were analysed and molecular weight was calculated using ASTRA 6 software (Wyatt). Values of dn/dc = 0.1926 ml g^-1^ and ε_280_ 3.198 ml (mg cm)^-1^ were used to calculate HHAT molecular mass. Protein conjugate analysis was performed with dn/dc = 0.1270 ml g^-1^ for OGNG (www.biorxiv.org/content/10.1101/2020.11.11.378570v1.full).

#### Bio-layer interferometry

All measurements were performed at 22 °C using SA biosensors in an Octet Red96e (both ForteBio). Biotinylated HHAT-Avi-1D4 was immobilised at 0.015-0.03 mg/mL in 10 mM HEPES, pH 7.5, 150 mM NaCl, 0.03 % DDM 40:1 CHS. For nanobody and megabody kinetic measurements, NB169, NB177, MB169 and MB177 were diluted into the same buffer at a 100 nM and a 1:3 dilution series created. For single-cycle kinetics, five 180 s association steps were each followed by 20 s dissociation steps and a final 600 s dissociation. For SHH binding tests, SHH samples were diluted into 10 mM HEPES, pH 7.5, 150 mM NaCl, 0.06 % DDM 40:1 CHS, 1 mM DTT, 5 mM CaCl2 with or without 5-10 mM nhPalm-CoA. Multi-cycle kinetic experiments were performed with association and dissociation steps of 300 s and a 1:1.5 SHH dilution series starting at 5 μM. Competition experiments were performed in a similar manner in a buffer containing 200 nM NB169 or MB177, with 300 s association and 600 s dissociation steps. IMP-1575 was added to a concentration of 10 μM where indicated. HHAT mutant studies were performed in the same manner with 0.015 mg/mL biotinylated HHAT mutants loaded and binding of 500 nM NB169 or 4 μM SHH. Loading was normalised to *wild-type* HHAT. Data were processed and analysed using ForteBio Data Analysis software (version 11.1). All kinetic data were double-referenced. Single-cycle kinetic data binding constants were calculated by the fitting of a 1:1 binding model for megabodies and nanobodies using MATLAB (R2021a). Multi-cycle kinetic data binding constants for SHH were calculated by the fitting of a one to two state binding model using TraceDrawer (Ridgeview Instruments).

#### UV/Vis spectroscopy

UV/Vis spectra were recorded in a range of 220-800 nm at room temperature in a 96-well UV-Star® Microplate (Greiner) using a CLARIOstar plate reader (BMG Labtech) with appropriate buffer blanks.

#### HHAT enzymatic activity using Acyl-cLIP

Biotinylated HHAT mutants and non-biotinylated HHAT mutants were analysed separately. For both groups, 5 µL of purified (mutant) protein sample was mixed with 1.67 µL of 4X Laemmli loading buffer (Bio-Rad 1610747), then 5 µL was loaded onto an SDS-PAGE gel (Either 12% isocratic (Bio-Rad 4561046) or 4-15 % gradient acrylamide (Bio-Rad 4561086DC)) and run at 200 V for 35 min. The gel was stained with InstantBlue Coomassie protein stain (Abcam ISB1L) and destained overnight. The gel was then imaged on an ImageQuant LAS 4000 (G.E. Electronics) and the obtained bands quantified using ImageJ (v. 1.53e) (Schindelin et al., 2012). From these band intensities, dilution factors to normalise each mutant to the lowest concentration sample (G287V) were calculated. All sample concentrations were then equalised using elution buffer (20 mM HEPES pH 7.5, 300 mM NaCl, 5% (v/v) glycerol, 0.03% (m/v) DDM 40:1 CHS, 0.5 mM 1D4 peptide). The equal concentration protein samples were split in two; one part was loaded onto an SDS-PAGE gel using the procedure described above and one part was used in Acyl-cLIP analysis of enzymatic activity as described below.

For the Acyl-cLIP assay, in a 384-well assay plate (Corning 3575) each well was filled with 4 µL reaction buffer (100 mM MES, pH 6.5, 20 mM NaCl, 1 mM DTT, 1 mM tris(2-carboxyethyl)phosphine (TCEP), 0.1% (w/v) BSA, filtered) and either 4 µL Palm-CoA solution (15 µM in reaction buffer) or another 4 µL of reaction buffer (negative control). Then, master mix buffer was prepared by adding 10% (v/v) DDM buffer (20 mM HEPES, pH 7.3, 350 mM NaCl, 5% (v/v) glycerol, 1% (w/v) DDM) to a 2 µM FAM-SHH peptide (synthesised in-house as previously described (Lanyon-Hogg et al., 2019)) solution in reaction buffer. The final master mix was then prepared for each (mutant) HHAT sample by adding 3 µL normalised protein solution to 100 µL of master mix buffer. 12 µL of HHAT containing master mix was then distributed to each well and the fluorescence polarisation (Excitation 480 nm, emission 535 nm) measured for 1 hour at 2 minute intervals on an EnVision Xcite 2104 (PerkinElmer). Enzymatic activity of each mutant was measured with (+ Palm-CoA) and without (-Palm-CoA) Palm-CoA and in quadruplicate.

The slope of the resulting curves was calculated for each individual well using Prism 9.1.2 (GraphPad) and each well was background subtracted by subtracting the average of its corresponding (-Palm-CoA) value. The background subtracted data were then normalised to wildtype activity and plotted in a bar chart using Prism.

